# Foraging on host synthesized metabolites enables the bacterial symbiont *Snodgrassella alvi* to colonize the honey bee gut

**DOI:** 10.1101/2023.01.23.524906

**Authors:** Andrew Quinn, Yassine El Chazli, Stéphane Escrig, Jean Daraspe, Nicolas Neuschwander, Aoife McNally, Christel Genoud, Anders Meibom, Philipp Engel

## Abstract

Dietary nutrients and microbial cross-feeding allow diverse bacteria to colonize the animal gut. Less is known about the role of host-derived nutrients in enabling gut bacterial colonization. We examined metabolic interactions within the evolutionary ancient symbiosis between the honey bee (*Apis mellifera*) and the core gut microbiota member *Snodgrassella alvi*. This Betaproteobacteria is incapable of metabolizing saccharides, yet colonizes the honey bee gut in the presence of only a sugar diet. Using comparative metabolomics, ^13^C tracers, and Nanoscale secondary ion mass spectrometry (NanoSIMS), we show *in vivo* that *S. alvi* grows on host-derived organic acids, including citrate, glycerate and 3-hydroxy-3-methylglutarate which are actively secreted by the host into the gut lumen. *S. alvi* additionally modulates tryptophan metabolism in the gut by converting kynurenine to anthranilate. These results suggest that *S. alvi* is adapted to a specific metabolic niche in the gut that depends on host-derived nutritional resources.

## Introduction

Gut bacteria and their hosts typically engage in mutualistic interactions. Metabolic exchange from the gut microbiota to the host is vital for uptake of essential nutrients, gut health and immune system function ^1^. In turn, bacteria profit from a stable niche environment and frequent supply of exogenous food. The role of host secreted metabolites that benefit bacteria in the gut is less well understood. Such metabolic exchange is difficult to identify due to the overwhelming contributions of the diet and microbial products towards gut metabolites.

Deciphering the extent to which host metabolite secretion drives microbial colonization of native symbionts is aided by a simple, tractable system in which diet and microbiota-derived metabolites can be tightly controlled. The Western honey bee (*Apis mellifera*) provides such a system: Its gut microbiota is broadly stable and composed of only 8-10 genera ^2,3^ many of which have longstanding evolutionary associations with their host that date back to the emergence of social bees >80 mya ^4–6^. Bacteria of these genera are culturable and can be inoculated individually or as defined communities into gnotobiotic bees ^7^. Furthermore, while the typical honey bee diet features a rich mixture of compounds found in nectar and pollen, bees can survive for extended periods on pure sugar water diets ^8^.

Most members of the bee gut microbiome are primary fermenters, possessing a broad range of carbohydrate degradation enzymes that enable them to utilize hemicellulose, pectin, starch, or glycosides found in pollen ^3,9^. A notable exception is the Betaproteobacteria *Snodgrassella alvi*. It colonizes the cuticular surfaces of the ileum in the hindgut and displays a markedly different metabolism ^10,11^. Lacking a functional glycolysis pathway, *S. alvi* profits from acids in the gut, generating energy from an aerobic TCA cycle, and biomass through gluconeogenesis. Fermentation by other microbial members has been proposed as the primary source of short chain fatty acids (SCFAs) consumed by *S. alvi*. In particular, bacteria of the Gammaproteobacterial genus *Gilliamella* are likely mutualistic partners for metabolic crosstalk, as they co-localize with *S. alvi* within biofilms attached to the cuticular surface in the ileum and share complementary metabolic capabilities ^10^. Experimental evidence from *in vitro* growth experiments bolstered this hypothesis, showing that *S. alvi* grows on the spent media of *Gilliamella,* while consuming numerous products of *Gilliamella’s* metabolism, such as succinate and pyruvate ^12^.

Although a strong case can be made for niche exploitation through bacterial cross-feeding, this hypothesis does not explain previous results in which *S. alvi* was able to mono-colonize bees fed diets of sugar water and pollen ^12^. To better understand the nutrient sources that *S. alvi* exploits, we provided bees a simple (sugar water) or complex (sugar water + pollen) diet and colonized them with *S. alvi* alone, or together with divergent strains of the genus *Gilliamella*. Surprisingly, we found that a simple sugar water diet was sufficient for *S. alvi* to colonize the honey bee gut. Subsequent metabolomics analysis indicated that host-derived carboxylic acids enable *S. alvi* colonization. We validated this hypothesis with a series of experiments to show that (i) these carboxylic acids are synthesized by the host, (ii) *S. alvi* utilizes them for growth, and (iii) the findings hold across a range of divergent *Snodgrassella* strains and species.

## Results

### *S. alvi* colonizes the honey bee gut in the presence of only sugar in the diet

We colonized microbiota-free (MF) honey bees with *S. alvi* (strain wkB2) or a mixture of *Gilliamella* strains (strains wkB1, ESL0169, ESL0182, ESL0297) or with both phylotypes together, and provided a diet composed of only sterile-filtered sugar water (sucrose) or sugar water and electron beam-sterilized polyfloral pollen for five days (**Fig. 1A**). Colonization levels of both phylotypes were assessed by qPCR and CFU plating five days post inoculation (**Fig. 1B**, **Fig. S1**). Surprisingly, *S. alvi* colonized at equivalent levels in the guts across all treatment groups, *i.e.*, sugar water in the diet was sufficient to enable *S. alvi* colonization and neither the addition of pollen to the diet nor co-colonization with *Gilliamella* significantly increased *S. alvi* loads (Wilcoxon rank sum test with *S. alvi* (sugar water) as reference group).

**Figure 1.**
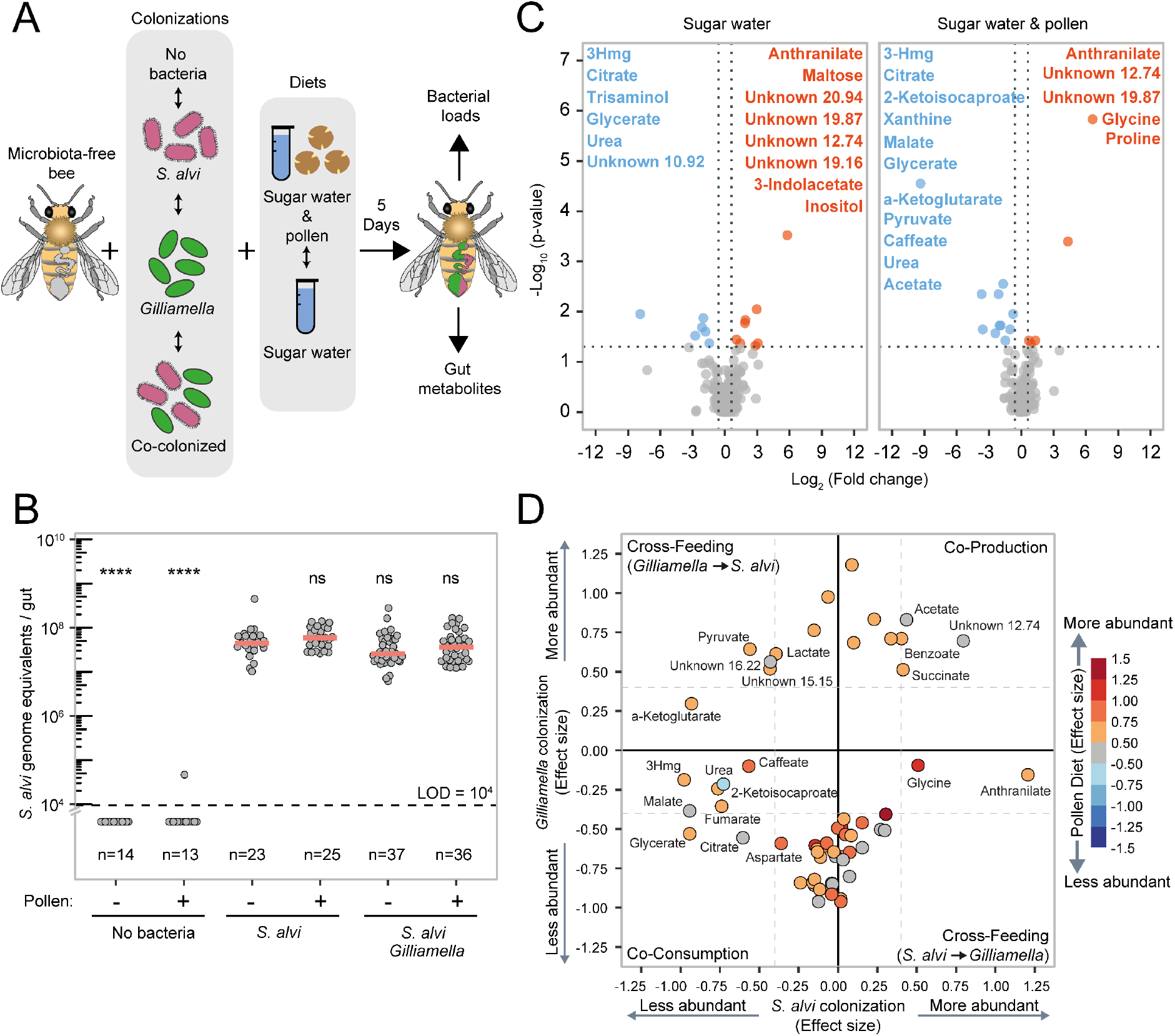
*S. alvi* colonizes bees fed with sugar water only and depletes host, pollen, and *Gilliamella*-derived metabolites. **(A)** Outline of the bee colonization experiment with four colonization treatments (no bacteria, *S. alvi*, *Gilliamella*, or co-colonized) and two dietary treatments (sterile sugar water and pollen, or sterile sugar water only). Five days after colonization gnotobiotic bees were sacrificed and the bacterial load and the metabolites in the gut quantified. **(B)** Number of *S. alvi* cells (number of bacterial genome equivalents) per bee gut determined by qPCR using *S. alvi*-specific 16S rRNA gene primers. Each value represents the number of cells in a single bee gut, red bars represent the median, horizontal black line represents qPCR limit of detection (LOD = < 10^4^). For each treatment, the presence/absence of pollen in the diet is indicated by +/-sign. Significance determined by Wilcoxon rank sum test with *S. alvi* (pollen -) as reference group, ns, non-significant; ****P < 0.001. (**C)** Significantly differentially abundant gut metabolites between *S. alvi* colonized and MF bees (i.e., ‘no bacteria’ treatment). Significantly depleted and produced metabolites are shown in blue and orange, respectively. Adjusted significance values were calculated using Wilcoxon rank sum test, and Benjamini-Hochberg correction, using the MF samples as a reference. **(D)** Results from mixed-linear modelling show the contributions (fixed effect sizes) of microbial colonization and diet towards z-score normalized metabolite abundances. Only metabolites significantly correlated with *S. alvi* colonization are named. Large values (+/-) indicate a strong correlation between a factor and a corresponding change in metabolite abundance. Values centered around zero (inside the dotted lines) are not significant. The effect size of pollen is indicated by the color of each metabolite, red colors indicate significantly positive correlation, while blue is a significant negative correlation. The four corners of the graph represent the four types of metabolic interactions between *S. alvi* and *Gilliamella* (from the top left: cross-feeding from *S. alvi* to *Gilliamella*, co-production by both bacteria, cross-feeding from *Gilliamella* to *S. alvi,* co-consumption). Carboxylic acids are grouped principally on the lower-left section of the plot indicating they are less abundant with *S. alvi* and *Gilliamella* colonization.

Importantly, qPCR analysis with universal bacterial and fungal primers showed no or very low levels of amplification in most samples, with the exception of gut samples containing pollen which resulted in relatively high background amplification with the universal bacterial primers as previously reported (**Fig. S1C, D**). Culturing of gut homogenates of MF bees on different media resulted in no microbial growth in any of the tested conditions (see methods). Hence, we can rule out that systematic contaminations with high levels of other microbes facilitated *S. alvi* colonization through cross-feeding in the mono-colonization treatment.

### *S. alvi* depletes organic acids in the honey bee gut

To search for putative *S. alvi* growth substrates originating from the host, the pollen diet, or *Gilliamella*, we next extracted metabolites from the mid-and hindgut for a subset of the colonized bees and analyzed them via GC-MS. The presence of pollen in the gut significantly increased the abundance of nearly half of the annotated metabolites (125/233) in MF bees (**Fig. S2**). Thus, we compared results between colonized and MF bees only within each dietary treatment. When we examined mono-colonization of *S. alvi* in bees fed with sugar water, we identified multiple carboxylic acids, including citrate, 3-hydroxy-3-methylglutarate (3Hmg), and glycerate that were significantly less abundant than in the MF controls **(Fig. 1C)**. These carboxylic acids, along with others (*e.g.*, 2-ketoisocaproate, alpha-ketoglutarate and malate) were also depleted in *S. alvi* mono-colonized bees that were fed pollen **(Fig. 1C)**, despite the substantial differences in the overall metabolite profiles between the two dietary conditions. In contrast, few metabolites were significantly more abundant in *S. alvi* colonized bees relative to MF bees, with only anthranilate, a product of tryptophan metabolism, accumulating in colonized bees of both dietary treatments (**Fig. 1C**).

We then fit each metabolite with a mixed linear model to quantify how strongly metabolite changes were influenced (*i.e.*, the fixed effect) by each of the three independent experimental variables (*i.e.*, the presence of *Gilliamella, S. alvi,* or pollen) across the eight experimental conditions, This cross-conditional analysis confirmed that colonization with *S. alvi* resulted in significantly (p-value < 0.05, see Supporting Data, “output.csv”) decreased abundances of carboxylic acids, particularly 3Hmg, citrate, malate, fumarate, and glycerate (**Fig. 1D**). We could also infer which metabolites were co-produced, co-consumed, or cross-fed (and in which direction) between the two bacteria (**Fig. 1D**). For example, we found evidence supporting our initial hypothesis of metabolic cross-feeding from *Gilliamella* to *S. alvi* in the form of lactate, pyruvate and other unknown compounds that were more abundant with *Gilliamella* and less abundant with *S. alvi* (**Fig. 1D**, **Fig. S3**). However, we found no evidence of reverse cross-feeding from *S. alvi* to *Gilliamella*. Instead, we found that both species compete for certain metabolites and act synergistically to synthesize others. Competition, or co-consumption, centered on the carboxylic acids, as colonization with *Gilliamella* also led to depletion of citrate and glycerate, and to a lesser extent, malate, and fumarate (**Fig. 1D**, **Fig. S3**). Cooperative synthesis revolved around four metabolites, acetate, succinate, benzoate, and unknown_12.74. Of these, only acetate was also more abundant in mono-colonization (*Gilliamella*) than in MF controls (**Fig. 1D**, **Fig. S3**). Finally, nearly all the metabolites depleted with *S. alvi* colonization were positively affected by pollen, except for citrate, glycerate, acetate, and two unknown compounds, which were unaffected, and urea, whose abundance was negatively correlated with pollen. Taken together, these results show that each of the tested variables impacts nutrient availability and metabolism of *S. alvi*, although metabolic cross-feeding and a nutrient rich diet do not lead to increased colonization levels in the gut.

### Host sugar catabolism provides *S. alvi* substrates *in vivo*

We next sought to measure host production of carboxylic acids in the guts of MF bees over the first six days post emergence and compare the abundances to what the bee could have extracted from the average amount of pollen consumed (24.3 ± 7.2mg/bee) as estimated based on data collected from the previous experiment (see methods, **Fig. S4** and Table S2). Moreover, in order to rule-out microbial or fungal contamination as a source of these compounds, we rigorously checked once more for live contamination by plating the guts of newly emerged (day zero) and six-day old MF bees on eight different media in three different growth environments (see methods), finding no evidence of live bacterial or fungal contamination (n=8). Even though the bees only consumed sugar water, we found that many gut metabolites, including glycosylamines, sugar alcohols and carboxylic acids, were significantly more abundant six days after emergence (**Fig. S5**). Focusing specifically on the compounds depleted in the presence of *S. alvi*, we found that citrate was most abundant in the gut, with its concentration increasing from 29 ± 14 μmol/mg at Day 0 to 73 ± 34 μmol/mg at Day 6 post emergence. Substantially less citrate, 3 ± 2 μmol/mg, was extracted from pollen (**Fig. 2A**). A similar trend was also found for 3Hmg, alpha-ketoglutarate, isocitrate and kynurenine. Of *S. alvi’s* putative substrates, only caffeate and lactate were more abundant in pollen than in the bee gut. The increasing abundance of many compounds over six days, as well as their generally low abundance in pollen suggests that active metabolism by the host substantially impacts the gut metabolome, i.e. easily digestible nutrients in the diet are absorbed and metabolized by the bee upstream of the hindgut, while downstream metabolic products are excreted into the hindgut and serve as substrates for *S. alvi*.

**Figure 2.**
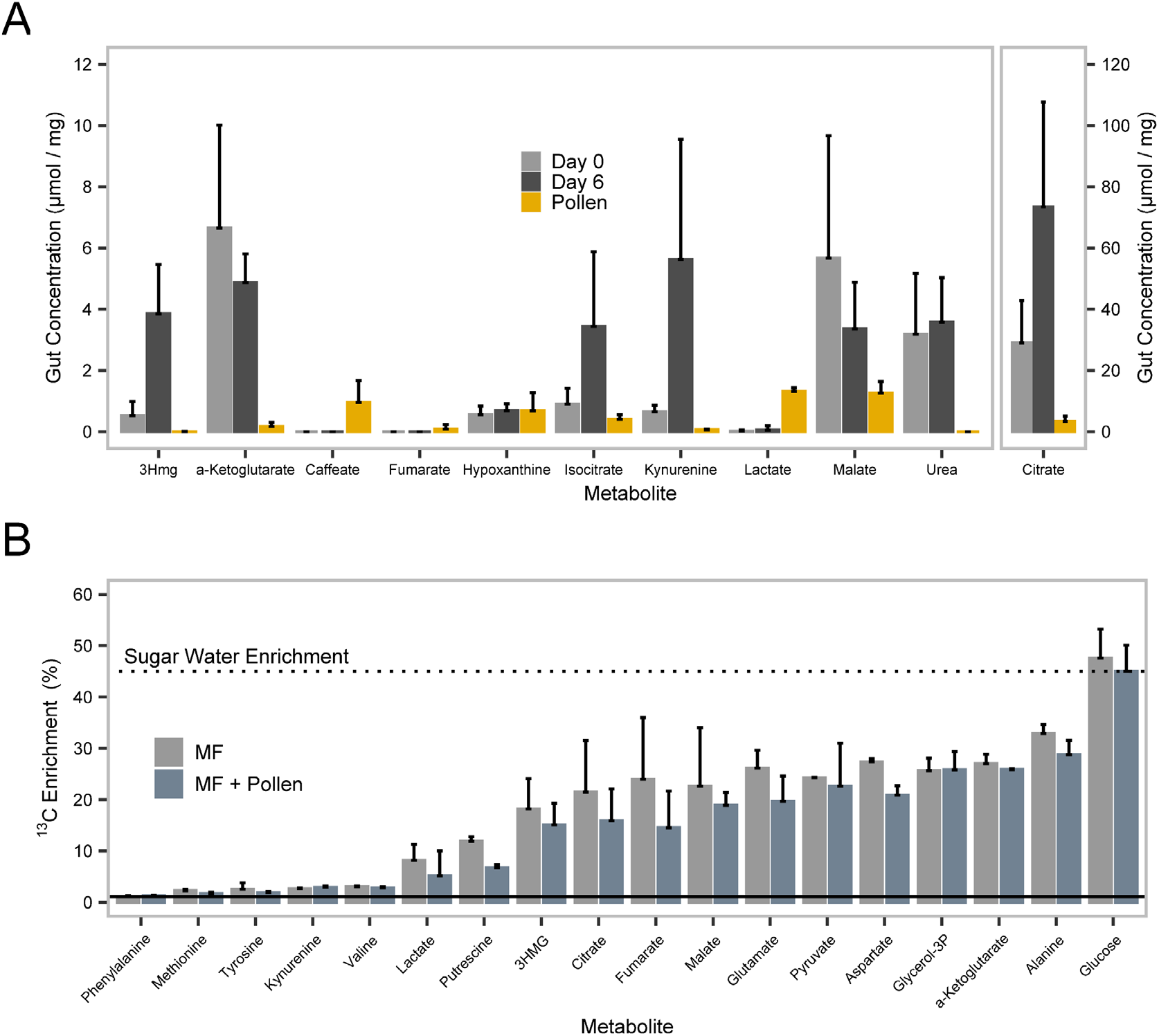
Gut metabolite abundances and isotope labelling reveals the host origin of *S. alvi* substrates. **(A)** Abundances of selected metabolites in the guts of MF bees increase from emergence (Day 0) to adult age (Day 6) (n=8). All metabolites except for caffeate and lactate were less abundant when extracted directly from an equivalent amount of pollen consumed per bee over six days (n=8). Quantitates from pollen extract were adjusted based on average total pollen consumption per bee (see methods). **(B)** The average (n=4) ^13^C metabolite enrichment (%) in bee guts shows that carboxylic acids and non-essential amino acids become highly labelled upon feeding ^13^C-glucose to MD bees. Dark and light bars indicate, respectively, absence and presence of pollen in the diet. The solid line denotes the natural ^13^C enrichment (1.108%), while the dashed line denotes ^13^C enrichment of sugar water solution fed to experimental bees.

To corroborate this hypothesis, we fed MF bees for six days with 45% of the dietary sucrose replaced by 100% U-^13^C_6_ Glucose. Half of the bees were also provided standard pollen to assess its contribution relative to simple sugars in the synthesis of gut metabolites. We then analyzed the resulting ^13^C isotopic enrichments in gut metabolites with a focus on those indicated as substrates for *S. alvi*. The carboxylic acids and non-essential amino acids were significantly ^13^C enriched in the bee gut. In contrast, other compounds, such as kynurenine and essential amino acids did not show any enrichment **(Fig. 2B)**. The lack of isotope labelling indicates that these compounds were not actively synthesized from glucose by adult bees during the first six days post-emergence. Instead, they were either leftover in the gut from the larval development stage or were acquired from the larval diet, such as in the case of essential amino acids and their catabolic products. As expected, the average ^13^C enrichment levels of most metabolites dropped in bees fed with pollen compared to those fed with sugar water only. The ^13^C dilution can occur directly from non-labelled metabolites in pollen, as well as indirectly from host metabolism of pollen substrates. However, the average ^13^C enrichment of carboxylic acids only dropped from 21 ± 6% to 17 ± 7%, which serves as further evidence that host metabolism of simple sugars rather than dietary consumption is the predominant source of carboxylic acids in the gut (**Fig. 2B).** We thus conclude from this analysis that the carboxylic acids utilized by *S. alvi* are mostly *de novo* synthesized from sugar metabolism of the host.

### NanoSIMS reveals transfer of host compounds to *S. alvi*

While our previous experiments suggested host nutrient foraging by *S. alvi*, the results did not provide direct evidence that *S. alvi* assimilates biomass from these compounds. Therefore, we probed the flow of metabolites from the bee to *S. alvi* using a “pulse-chase” ^13^C isotope labelling experiment (ILE), and then measured the enrichment within *S. alvi* cells and surrounding host tissue using nanoscale secondary ion mass spectrometry (NanoSIMS) complemented with measurements of metabolite enrichments using GC-MS (**Fig. 3, Dataset S1**). To do so, we enriched newly emerged MF bees in ^13^C by feeding them 100% U-^13^C_6_ glucose for four days after emergence. We then inoculated them with *S. alvi* and waited one day before switching their diet to naturally abundant, 98.9% ^12^C-Glucose (**Fig. 3A**). This ensured that the largest *S. alvi* population increase, between 24-and 48-hours post-colonization, occurred without a dietary ^13^C source, but still in a highly ^13^C labelled environment **(Fig. 3B)**.

**Figure 3.**
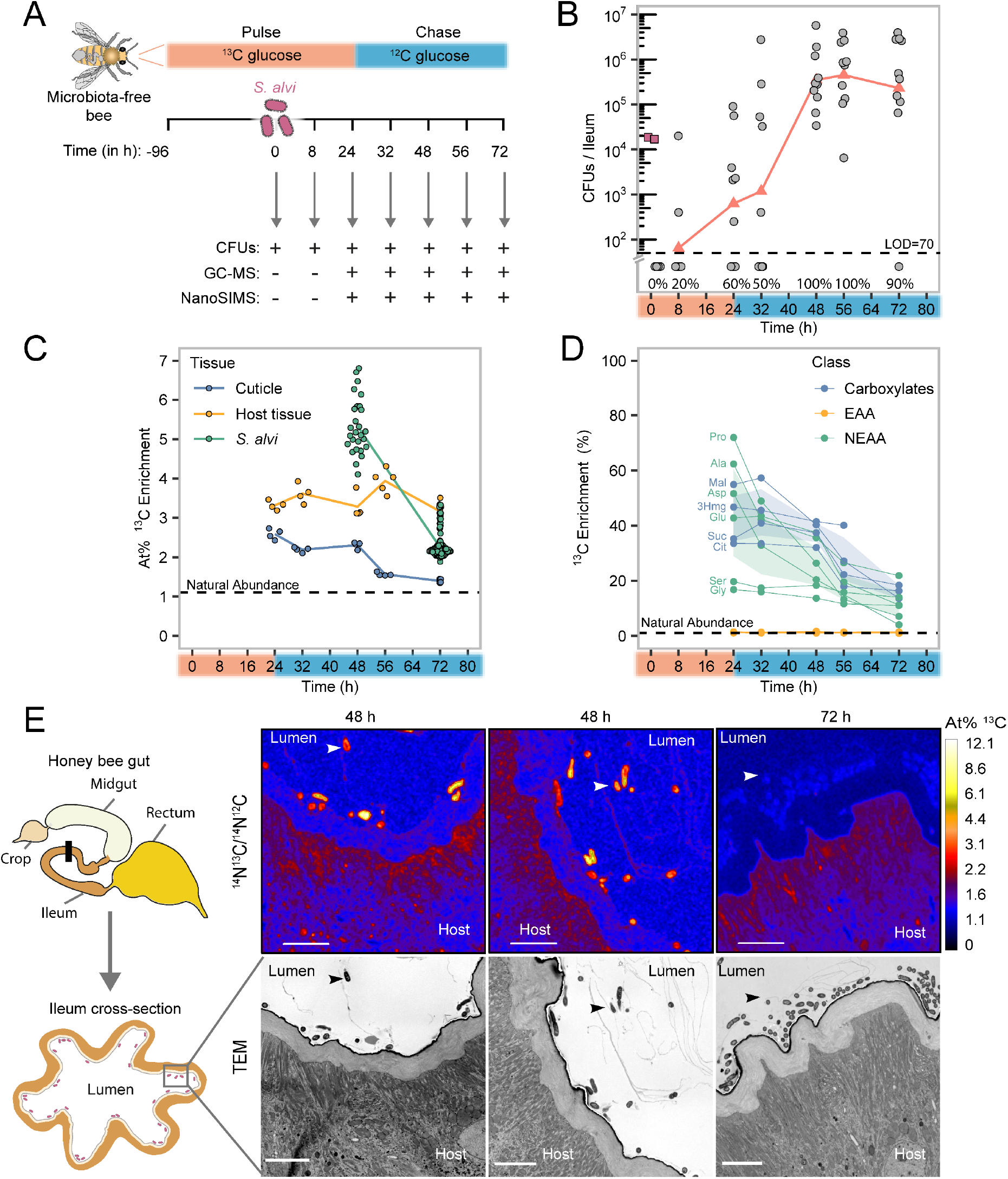
*S. alvi* builds biomass from substrates derived from the glucose metabolism of the host. **(A)** Experimental design of the pulse-chase isotope labelling gnotobiotic bee experiment. Newly emerged MF bees were fed 100% ^13^C glucose for four days (i.e., 96 hours), colonized with a defined quantity of *S. alvi* and then the ^13^C glucose was replaced by ^12^C glucose 24 hours after colonization. The timeline indicates when bees were sampled for CFU plating, GC-MS and NanoSIMS analysis relative to the timepoint of colonization (i.e. 0h). **(B)** Colonization levels of *S. alvi* in the ileum until 72 hours post inoculation show the initial strong colonization bottleneck and subsequent exponential increase in cell numbers. Colonization success is indicated as percentage of bees with detectable CFUs. Pink squares indicate the inoculum of *S. alvi* (OD= 0.1). The limit of detection (LOD) of 70 CFUs per ileum is indicated with the dashed line. **(C)** The average At% ^13^C enrichments of *S. alvi* cells decreases rapidly, while the enrichment of host cells and epithelium remains constant over 72 hours in ileum cross-sections imaged with NanoSIMS. Each data point represents the mean of regions of interest (ROI); bacterial ROIs consist of single bacterial cells, while epithelium and host cells ROIs represent the total area encompassing epithelium or host cell tissue in each image. ROI raw counts and calculated enrichments are listed in **Dataset S1**. The dashed line at 1.1 % indicates the natural ^13^C abundance calculated from NanoSIMS images of ^12^C control bees. **(D)** ^13^C enrichment of metabolites in the gut steadily decreases in the ^12^C chase phase. Values represent the average ^13^C enrichment of a given metabolite across the bees sampled at that timepoint. Average values and colors are shown by metabolite class: carboxylic acids (violet), non-essential amino acids (NEAA, green), essential amino acids (EAA, brown). Measured metabolites are listed in **Dataset S1**. The dotted line at 1.1 % indicates the natural ^13^C abundance. **(E)** NanoSIMS and corresponding TEM images of 2 different ^13^C labeled bees at 48 and 72 hours after inoculation. White arrows indicate *S. alvi* cells. At% ^13^C represents percentage of ^13^C atoms; the natural ^13^C abundance is ∼1.1 at%. Scale bars = 5 μM. Schematic drawing of the honey bee gut shows the region in the ileum (black bar) where cross-sections were taken and which regions of the cross-sections were imaged.

Accordingly, we measured rapid ^13^C labelling turnover in *S. alvi* in 18 images coming from two bees (sampled at 48-and 72-hours post-inoculation) in which we could detect bacterial cells **(Fig. 3C, E, Dataset S2)**. The ^13^C enrichment was substantially higher in *S. alvi* cells than in the adjacent host ileum tissue 48 hours after inoculation, but then dropped below the levels of host tissue 72 hours after inoculation (At% = start: 5.56, end: 2.25) **(Fig. 3C)**. In contrast, the ileum tissue displayed a constant ^13^C enrichment over the course of the experiment (At% = start: 3.31, end: 3.16), whereas the cuticle lining of the gut epithelium was comparatively less enriched initially and continued dropping to almost natural ^13^C enrichment levels after 72 hours (At% = start: 2.58, end: 1.39) **(Fig. 3C)**.

The ^13^C enrichment of carboxylic acids and host synthesized (non-essential) amino acids in the gut dynamically shifted in response to the ^13^C/^12^C switch (**Fig. 3D, Dataset S1)**. Initially, they were highly ^13^C enriched (38 ± 14.7%), but the labelling fell steadily to approximately half the original level (14 ± 5.8%) within 48 hours (**Fig. 3D)**. As a control, we found that essential amino acids (non-host synthesized) were not enriched above the level of natural ^13^C abundance (1.11%) throughout the experiment. Thus, these results show that products of host glucose metabolism are actively used by *S. alvi* to build its biomass during early colonization.

### *In vitro* growth assays confirm active metabolism by *S. alvi*

We next tested whether the putative host synthesized substrates were sufficient for growth of *S. alvi* as sole carbon sources in a chemically defined liquid medium “Bee9”, which we derived from standard M9 media (**Fig. 4A**, **Dataset S3**). We utilized Bee9, after finding that *S. alvi* consumes amino acids present in standard M9 previously used to grow this bacterium ^12^. Only three metabolites (citrate, isocitrate, and malate) supported growth as sole carbon sources in this condition. However, the addition of 3Hmg, fumarate, succinate, gamma-aminobutyrate (Gaba), kynurenine, or urea to Bee9 + citrate significantly improved the maximal growth rate of *S. alvi* relative to Bee9 + citrate alone (**Fig. 4B**). In particular, 3Hmg and Gaba had dramatic effects on growth rates, increasing them 3.52 ± 0.10 and 2.76 ± 0.04-fold, respectively.

**Figure 4.**
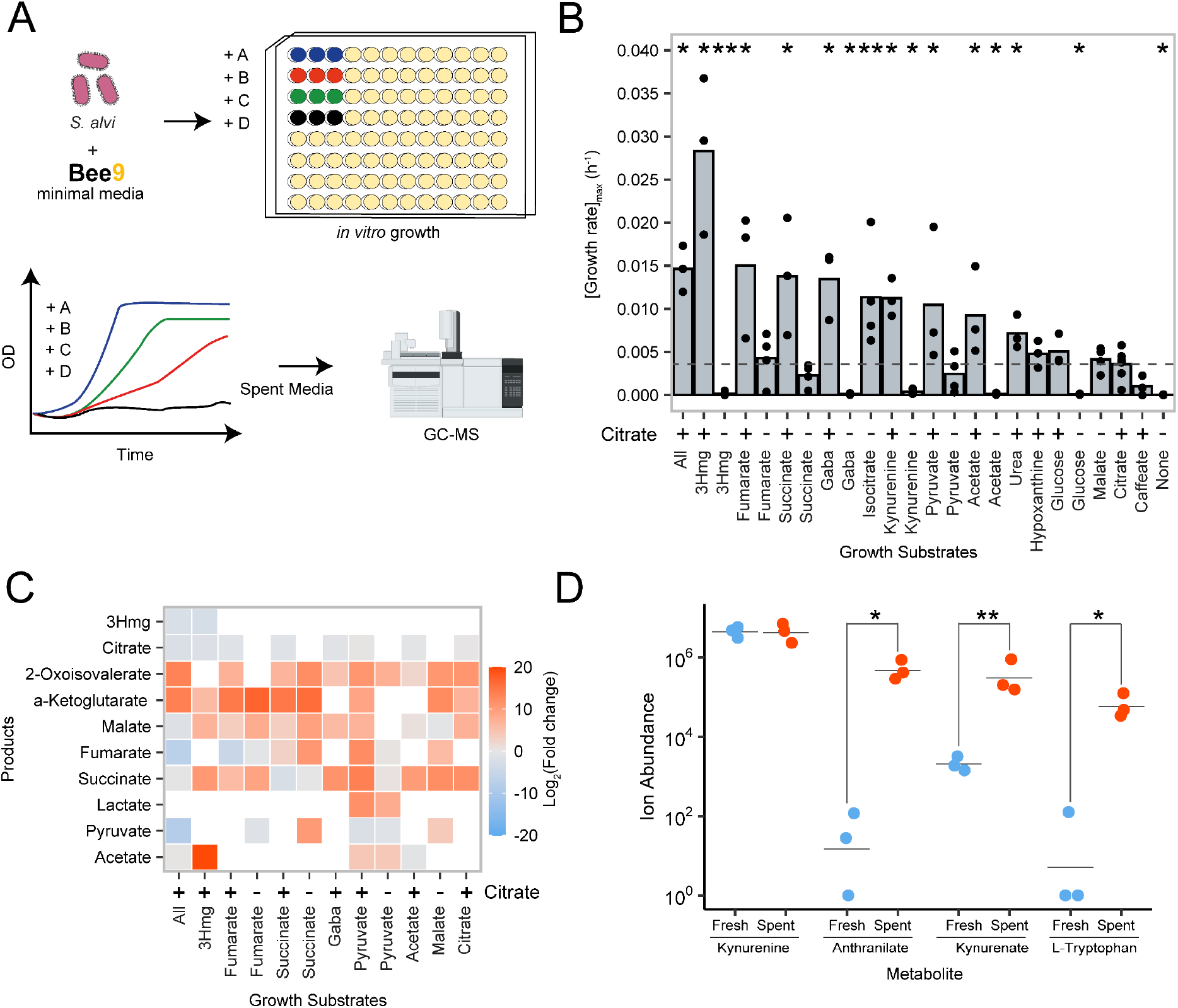
Host synthesized compounds enhance *S. alvi* growth *in vitro*. **(A)** Schematic diagram depicting growth and spent supernatant analysis. The “Bee9” media was derived from M9 base media (**Dataset S3**). **(B)** The maximum growth rate across single carbon sources with (+) or without (-) citrate added to the medium, reveals that growth on citrate + 3Hmg was faster than for all other conditions, including “All”, where all substrates were pooled together. The average value (bars) was calculated from the individual biological replicates (dots). Adjusted significance values were calculated using Wilcoxon rank sum test, and Benjamini-Hochberg correction, using the growth on citrate as the reference state (dotted line). **(C)** Abundances of carboxylic acids in the spent supernatant, relative to growth on citrate alone, varies with growth substrates. While some carboxylic acids including a-ketoglutarate and malate are produced across most conditions, acetate is a growth substrate, but only produced with either pyruvate or 3Hmg in the media. **(D)** Addition of kynurenine to the media leads to the production of anthranilate, kynurenic acid, and L-tryptophan. Significant changes between fresh and spent media were calculated using a paired t-test. (* P < 0.05, ** < 0.01, *** < 0.001, **** < 0.0001).

Analysis of the spent media revealed that all tested metabolites, except for the glucose negative control, were depleted by *S. alvi* (**Fig. S6**). This further confirmed that the identified host-derived organic acids are growth substrates of *S. alvi.* Unlike the results from our colonization experiments, *S. alvi* produced numerous carboxylic acids, which varied based on the carbon source. For example, acetate was a metabolic substrate, but it was also produced in the presence of pyruvate or 3Hmg. Meanwhile, alpha-ketoglutarate, 2-oxoisovalerate, malate and succinate were produced across most growth conditions (**Fig. 4C**). Consistent with the *in vivo* metabolomics, we detected production of anthranilate when kynurenine was added to the media. Additionally, we also measured the production of kynurenic acid and tryptophan (**Fig. 4D).** *S. alvi* wkB2 encodes a putative kynureninase gene in its genome (SALWKB2_0716, Genbank genome: CP007446 ^10^) that is likely responsible for the conversion of kynurenine into anthranilate. Interestingly, this gene seems to have been acquired by horizontal gene transfer as indicated by a phylogenetic analysis **(Fig. S7)**: apart from other *Snodgrassella* strains, the closest sequences of the kynureninase gene were not found in other Neisseriaceae, but in more distantly related Betaproteobacteria (genera *Pusilimonas* and *Alcaligenes*) and in Gammaproteobacteria (genera *Ignatzschineria* and *Acinetobacter*).

### Colonization without an exogenous nutrient source is conserved across strains of the genus *Snodgrassella*

As a final step, we examined whether our results were unique to *S. alvi* wkB2 type strain, or were more generally applicable to the genus, by inoculating bees with five divergent strains of *Snodgrassella*, three native to honey bees and two native to bumble bees. We also lengthened the colonization experiment from five to ten days to account for a potential delay in colonization of the honey bee gut by non-native strains. All five strains successfully colonized bees fed only sugar water. However, the efficiency and extent of colonization varied between strains. Strains of *Snodgrassella* native to honey bees colonized consistently, while non-native strains sometimes failed to colonize (Aggregate Colonization Success: 100% vs. 70%; Fisher’s Exact test: p < 0.001) (**Fig. 5A**). The gut metabolomic comparison between successfully colonized versus MF bees was qualitatively similar to our initial results. Carboxylic acids, such as 3Hmg, malate, fumarate, succinate, isocitrate, and glycerate were significantly less abundant in colonized guts vs MF controls across three or more of the *S. alvi* strains tested. The levels of purine and amino acid precursors hypoxanthine and urea were also depleted in the guts of colonized bees, while anthranilate again accumulated in all colonized bees (**Fig. 5B, Fig. S8**). These results show that the utilization of host-derived carboxylic acids is a conserved phenomenon in the genus *Snodgrassella* and facilitates gut colonization independent of bacterial cross-feeding or diet-derived nutrients.

**Figure 5.**
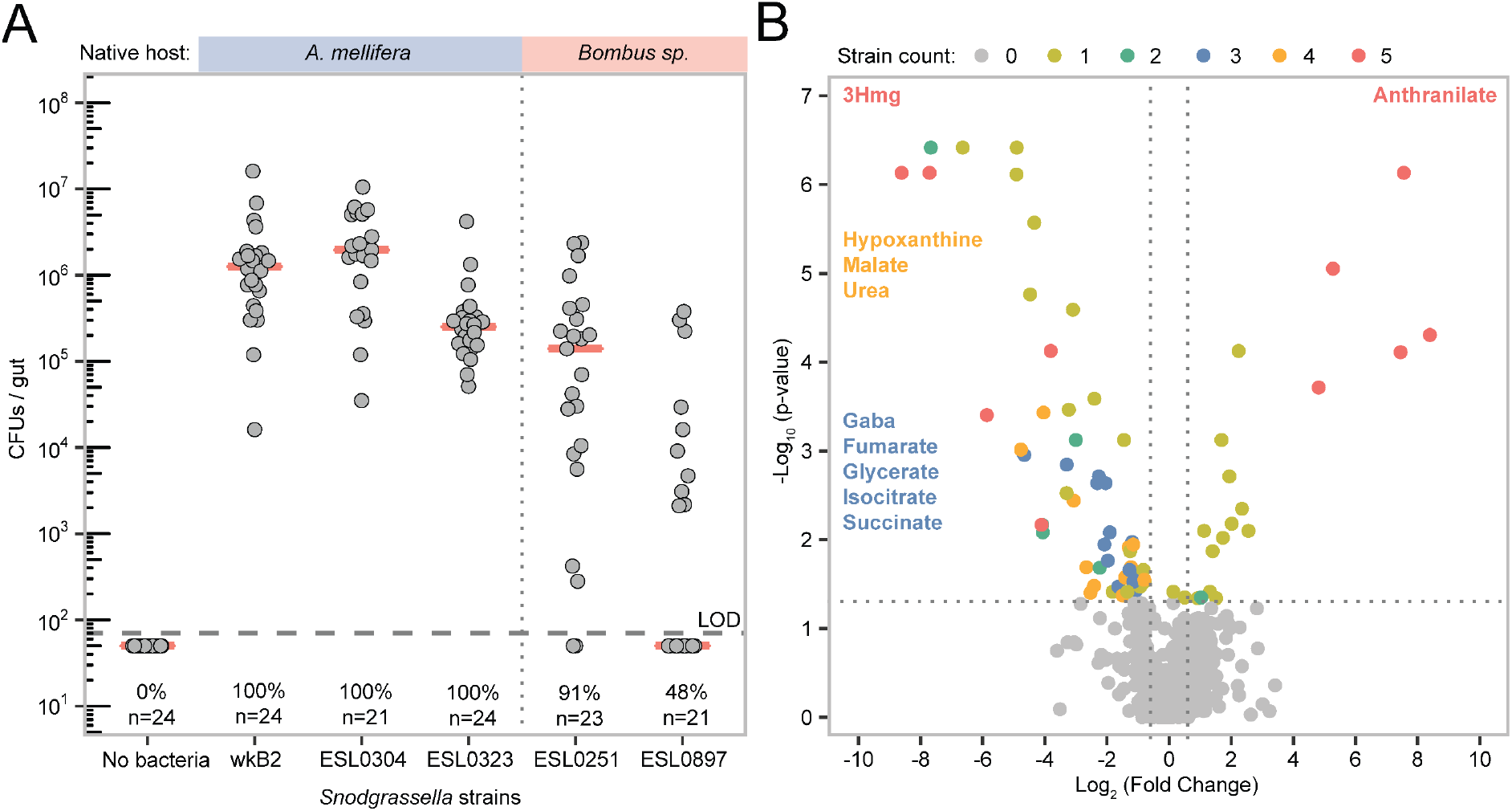
Gut colonization in the presence of a sugar-restricted diet and depletion of host-derived metabolites is conserved across divergent *Snodgrassella* strains. **(A)** Colonization levels of divergent *Snodgrassella* strains in monocolonized bees. Pink bars denote the median of CFUs/gut for each group. The LOD=70 colonies/gut is shown with a horizontal dashed line. The top bar indicates *Snodgrassella* strains native to *A. mellifera* versus *Bombus sp.* Colonization success (%) is shown for each strain, defined as guts with detectable CFU counts. (**B**) Volcano plot of gut metabolites showing the similarities in metabolite changes from MF vs colonized bees across strains. Each metabolite is plotted five times (once per *Snodgrassella* strain), and color coded by the number of strains in which it is significantly differentially abundant vs MF bees control. Only 3Hmg and anthranilate were significantly depleted or enriched by all five strains, while four of the strains depleted hypoxanthine, malate, and urea, and three of the five depleted Gaba, fumarate, glycerate, isocitrate and succinate.

## Discussion

Host-derived metabolites are gaining appreciation for their importance in facilitating extracellular microbial colonization across widely disparate animal models. In some cases, bacteria graze on the chitinous lining of the murine gut or of light organ in squid ^13–20^. In other instances, bacteria profit from small molecules secreted into the lumen. Commensal species can utilize lactate, 3-hydroxybutyrate and urea in the murine gut, while pathogenic *Salmonella* species utilize aspartate, malate, lactate or succinate to invade the gut ^21–24^. Our study demonstrates that this phenomenon is conserved across widely disparate animal hosts and that these host-derived metabolites can represent the major carbon source for certain gut bacteria facilitating colonization and growth, independent of the diet or cross-feeding.

We took advantage of the bee as a model host to drastically minimize confounding factors, colonizing bees with a single bacterium, *S. alvi,* while restricting the host to a sole dietary substrate that is undigestible for this bacterium. We then utilized ^13^C-glucose labelling to show that organic acids measured in the gut are synthesized by the host from dietary sugars which are then assimilated by *S. alvi* cells during gut colonization. The ^13^C enrichment of *S. alvi* was initially higher than in the surrounding host tissue, but then dropped after the host diet was switched to ^12^C-glucose. These dynamics are consistent with the hypothesis that *S. alvi* utilizes host metabolites that are derived from simple carbohydrate catabolism in the bee diet, rather than from grazing on components of the host epithelium lining in the gut (**Fig. 6**). Finally, we demonstrated that these organic acids sustain growth of *S. alvi in vitro*.

**Figure 6.**
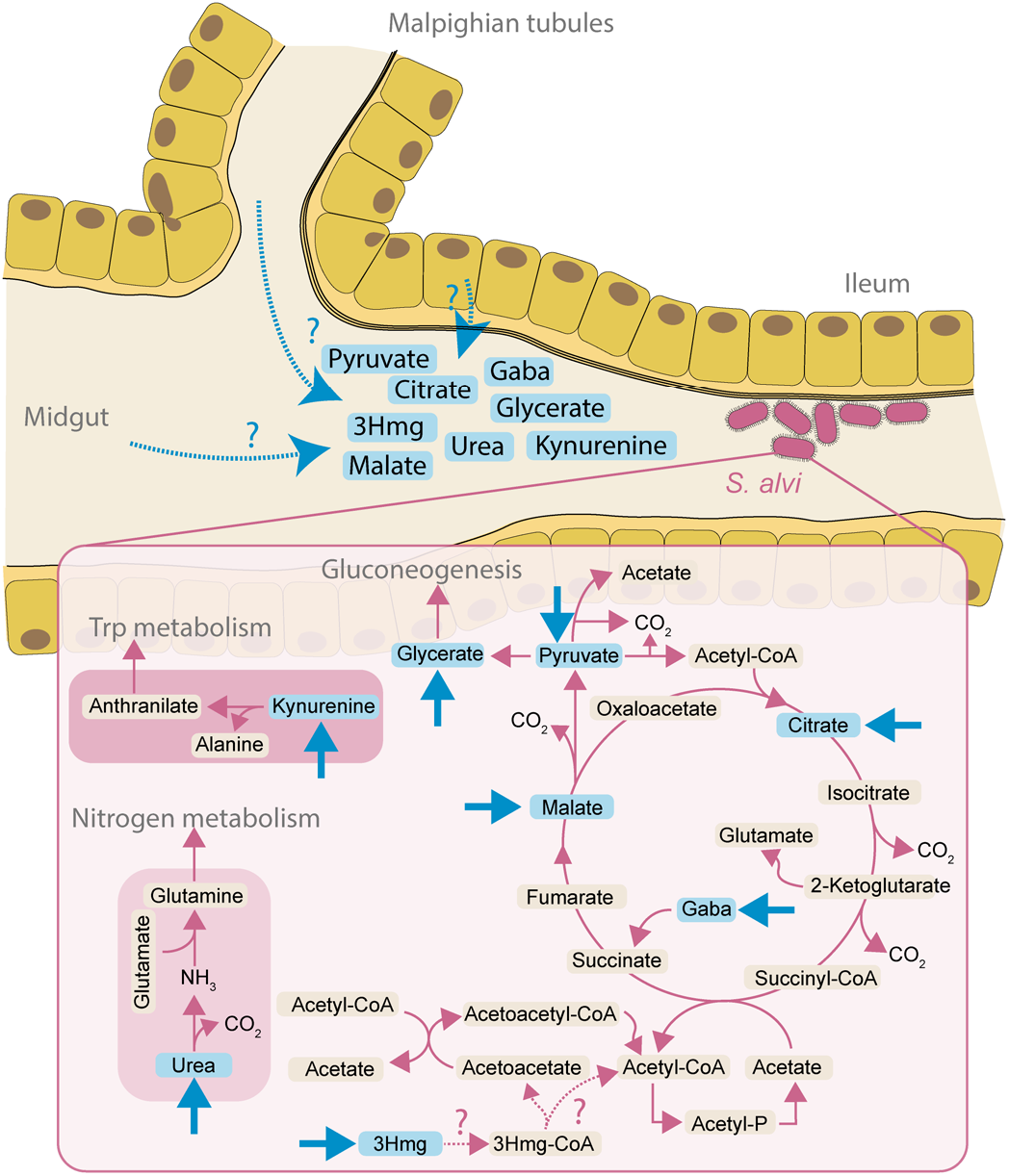
Schematic model of *S. alvi* metabolism of host-derived compounds in the ileum. Host metabolites (blue) enter the ileum through the epithelial cells, or through the malpighian tubules upstream of the ileum. *S. alvi* primarily utilizes the TCA cycle to generate energy, then synthesizing biomass through gluconeogenic reactions. 3Hmg provides a source of acetate to fuel the TCA cycle. However, the reactions synthesizing the first two steps (dotted pink arrows) have not been annotated in the genome. Additionally, host synthesized urea is a source of nitrogen, and kynurenine is converted to anthranilate as part of tryptophan metabolism. Blue arrows indicate the entry of substrates provided by the host metabolism.

Our results provide new context to the substantial body of research on bacterial colonization of the bee gut. The consistent depletion of host-derived acids linked to the TCA cycle complements findings that these metabolites are depleted in bees colonized with individual microbes including *S. alvi*, *Gilliamella, Lactobacillus* Firm-5 and *Bartonella apis*, as well as when colonized with a full microbial community^12,25,26^. Thus, foraging on host-derived compounds, although not always essential, may be widespread in the native bee gut community. Yet, in the specific case of *S. alvi*, these host-derived nutrients seem to be key for colonization. This is supported by a previous study which found that all genes of the TCA cycle and multiple genes associated with organic acid and ketone body degradation, provide a strong fitness advantage to *S. alvi* to grow in the bee gut ^27^.

Strikingly, two major metabolic functions conserved across all five strains of *S. alvi* (3Hmg consumption and anthranilate production), were associated with unique features in the metabolic pathways of *S. alvi*, providing evidence for adaptation to a specialized host niche. *S. alvi* possesses a non-canonical TCA cycle (**Fig. 6**); lacking a glyoxylate shunt it is unable to grow on acetate alone, but the substitution of acetate:succinate CoA-transferase for the canonical succinyl-CoA synthetase means that acetate, rather than acetyl-CoA, is a key driver of the TCA cycle ^28^. Interestingly, we found that 3Hmg consumption results in the production of acetate when *S. alvi* is grown on 3Hmg *in vitro*. Therefore, we postulate that host-derived 3Hmg enhances growth of *S. alvi* via the production of acetate, which in turn, increases the flux through the TCA cycle (**Fig. 6**). 3Hmg is an intermediate of the host’s isoprenoid biosynthesis, leucine degradation and ketone body metabolism, but the reason why this metabolite is released into the gut remains elusive.

The second unique metabolic feature of *S. alvi*, the production of anthranilate, likely depends on the enzymatic activity of a kynureninase gene identified in the genome of *S. alvi* converting host-derived kynurenine into anthranilate (**Fig. 6**). Intriguingly, this is the only gene annotated in the tryptophan degradation pathway of *S. alvi*; it is highly conserved in the entire genus and has likely been acquired by horizontal gene transfer, suggesting that it encodes a conserved function that is specific to the symbiosis between *Snodgrassella* and its bee hosts.

The release of relatively valuable metabolites into the gut could be a way for the host to control community assembly and facilitate the colonization of particularly beneficial gut symbionts ^29^. While the full gut microbiota has been shown to carry out several important functions for the host, the specific role of *S. alvi* has remained elusive. It is possible that its metabolic activities, such as the conversion of kynurenine into anthranilate, has a positive effect on the host, such as protection against pathogen invasion ^30,31^. Kynurenine is important during the larval stage of bee development, but it is also associated with neuronal defects (*e.g.*, hyperactivity and motor disfunction) in insects and vertebrates ^32–34^. In contrast, anthranilate is an important precursor of the essential amino acid tryptophan and several neurotransmitters such as serotonin, tryptamine and various indole derivatives, all of which are highly beneficial to bees ^35^. Intriguingly, indoles synthesized from tryptophan by *Lactobacillus* Firm-5 species in the gut were recently linked to enhanced memory and learning in honey bees ^36^. Finally, we also note that the metabolism of urea by *S. alvi* could have a beneficial effect on the host. Urea constitutes a major waste product in mammals and insects. In turtle ants, ancient, specialized gut bacteria are able to recycle large amounts of nitrogen from urea into both essential and non-essential amino acids ^37^. We found that urea is more abundant in the bee gut when exogenous nitrogen was absent from the diet, indicative of higher metabolic turnover and waste, which *S. alvi* could alleviate through urea fixation.

Our results and analysis fit within the context of several limitations. We examined a portion of the total gut metabolome that is amenable to analysis with gas chromatography. Further compounds of interest may be detected using complimentary analytical capabilities. While we have shown that host-derived compounds can sustain growth of *S. alvi*, the tested conditions did not reflect the natural state of the bee gut with a full microbial community and a metabolically rich diet that alters between bee bread, pollen, honey and flower nectar. Therefore, we cannot exclude that dietary or microbially-derived metabolites contribute to the growth of *S. alvi* in the native gut. In fact, we found *in vivo* evidence for cross-feeding of lactate and pyruvate from *Gilliamella*, as well as for pollen as an indirect source of key organic acids in the gut. However, these nutrient sources seem to play a minor role relative to the host, as the colonization levels of *S. alvi* did not differ significantly when pollen and/or *Gilliamella* were present. This is consistent with a previous study, which showed that the total abundance of *S. alvi*, in contrast to most other community members, does not change between nurse and forager bees that have different dietary preferences, nor does it change when pollen is removed from the diet of fully colonized bees under laboratory conditions ^38^.

How host synthesized organic acids reach the gut lumen remains elusive (**Fig. 6**). While leakage from host cells across the epithelial barrier is possible, transport via the malpighian tubules upstream of the ileum seems more likely, as they excrete nitrogenous waste and other metabolites into the gut while also regulating osmotic pressure in the hemolymph ^39,40^. Future work may resolve this question through careful dissection and metabolomic analysis of the malpighian tubules and the ileum, possibly with the assistance of labeled compounds injected into the bee thorax.

## Methods

### Bacterial culturing

**Table S1** lists all strains used in this study. Strains were isolated by directly plating gut homogenates on brain heart infusion agar (BHIA), or by first culturing them in Insectagro™ (Gibco) liquid media under microaerophilic conditions for 48 hours before colonies were streaked on plates. Species identity was confirmed by 16S rRNA gene sequencing. Strains were grown on liquid Insectagro™ media for 18 hours and then diluted in Phosphate buffer saline (PBS) and 25% glycerol at OD_600_ =1. Solutions were kept frozen at −80 °C until just before colonization, when they were thawed on ice, diluted 10-fold with PBS and mixed 1:1 with sterile sugar water (1kg/L of sucrose in water).

### Experimental colonization of gnotobiotic bees

All honey bees were collected from hives at the apiary of the University of Lausanne. A different hive was selected for each colonization replicate. Microbiota-free bees were raised according to established protocols ^41^. Briefly, frames containing capped brood were taken from the hive and washed quickly with a towel soaked in 3% bleach. Pupae of the appropriate age were then removed from the frames and placed into sterile emergence boxes containing 0.2 μm filter sterilized sucrose (50% w/w) sugar water. The boxes were placed in an incubator at 35 °C for 2 days, until the adult bees emerged. They were then randomly transferred to separate, sterile cages assembled from plastic drinking cups and Petri dishes.

Bees were colonized by hand with 5 μL of bacterial solution, or corresponding blank control, after starving them of sugar water for 1-2 hours, stunning them on ice for 5 minutes, and transferring them with tweezers to individual microcentrifuge tubes that were cut to enable insertion of a pipette tip. Sugar water and, where specified, polyfloral pollen (Bircher Blütenprodukte) sterilized via electron beam irradiation (Studer Cables AG) were provided to cages *ad libitum* throughout each experiment. Mono-and co-colonization of bees with *S. alvi* wkB2 and the four *Gilliamella* strains were performed five times, using different hives for each treatment. Not all conditions were run in each experiment due to technical difficulties preparing the treatment groups, and the GC-MS analysis was only carried out for a subset of the samples (see **Table S2** for details). The colonization experiment with the divergent *Snodgrassella* strains isolated from honey bees and bumble bees was carried out twice (see **Table S3** for details).

At the end of each experiment, bees were anesthetized with CO_2_, placed on ice, and dissected with sterile forceps. Whole guts, minus the crop, were homogenized in a bead beater (FastPrep-24, MP) in tubes containing PBS and glass beads. The samples were each immediately divided into three aliquots. One was plated to count Colony Forming Units (CFUs) on BHIA as well as for contaminants on Nutrient agar (NA) and De Man, Rogosa and Sharpe agar (MRSA). Another aliquot of 500 μL was centrifuged for 15 minutes at 4 °C and 20,000 g^-1^ and the liquid supernatant snap frozen in liquid nitrogen and stored at −80 °C until metabolomic extraction. The third aliquot was frozen at −80 °C until DNA extraction.

### Validation of bee sterility

To rule out contamination in the guts of newly emerged bees, two bees from each emergence box in each experiment were sacrificed and 10 μL of their homogenized guts plated on BHIA (5% CO_2_), NA (aerobic) and MRSA (anaerobic) at 35°C to check for sterility prior to colonizing the remaining bees.

In the event of contamination detected in newly emerged bees, all bees from that emergence box were discarded from the experiment. The same contamination checks were also performed on all analyzed bees at the end of each experiment. Additional sterility checks were performed in the experiment where we compared gut metabolite levels in newly emerged and six-day old MF bees. Here we also plated eight homogenized guts on Potato dextrose agar (PDA, aerobic), Lysogeny broth agar (LBA, aerobic), Tryptic soy agar (TSA, 5% CO_2_), Columbia agar + 5% sheep blood (CBA+5%SB, 5% CO_2_ & anaerobic), and Tryptone yeast extract glucose agar (TYG, anaerobic).

### Quantification of pollen consumption

Both whole bees and dissected bee guts were weighed to assess treatment effects and quantify pollen consumption. Whole bee and wet gut weights were both significantly higher in bees provided pollen than in those provided sugar water only, while colonization did not significantly affect weight (**Fig. S4**). (Linear mixed effects models fitted by ML with nested, random cage effects: [Bee weight, n=271, Diet: F_(1,262)_=176.1, p<.001; *S. alvi*: F_(1,262)_=0.1, p=.769; *Gilliamella*: F_(1,262)_=1.0, p=.311; *S. alvi:Gilliamella*: F_(1,262)_=0.7, p=.388]; [Gut weight, n=271, Diet: F_(1,262)_=220.0, p<.001; *S. alvi*: F_(1,262)_=0.3, p=.595; *Gilliamella*: F_(1,262)_=0.02, p=.880; *S. alvi:Gilliamella*: F_(1,262)_=2.3, p=.131]. We also weighed filled pollen troughs at the beginning and end of the experiment and calculated the amount of pollen consumed per bee. Unsurprisingly, the mean difference in gut weights between dietary groups (22.3 ± 18.3 mg/bee) closely matched the measured amount of pollen consumed by the bees across colonization treatments (24.3 ± 7.2 mg/bee) (**Table S2**). We then calculated the bee body weight (bee weight – gut weight) to determine if diet or colonization led to actual tissue weight gain in bees. While we found a slight increase in body weight from pollen (95% CI [0.69 mg, 7.51 mg]), we cannot rule out that this was in fact due to variable amounts of pollen present in the crop, which was not removed with the hindgut (Gut weight, n=271, Diet: F_(1,262)_=5.2, p=.023; *S. alvi*: F_(1,262)_=1.4, p=.240; *Gilliamella*: F_(1,262)_=0.6, p=.442; *S. alvi:Gilliamella*: F_(1,262)_=3.8, p=.052).

### Isotope tracing experiments

To assess the ^13^C metabolite enrichment in the bee gut, MF bees were provided a 45:65 ratio of 99% uniformly labelled U-^13^C_6_ glucose (Cambridge Isotope Laboratory) and naturally abundant sucrose (Sigma Aldrich) in the first experiment. For the NanoSIMS time-course colonization experiment, newly emerged MF bees were divided into two cages and provided pure 99% U-^13^C_6_ glucose (treatment) or naturally abundant (*i.e.* ∼98.9% ^12^C) -glucose (control) in water for five days (pulse) before switching both cages to a naturally abundant glucose diet (chase). On the morning of the fourth day all bees were starved for 1.5 hours and colonized with *S. alvi* wkB2. The ^13^C glucose solution was replaced with a ^12^C glucose solution 24 hours later. Bees were harvested at 0, 8, 24, 32, 48, 56, 72 hours after inoculation. This overlap of colonization and ^13^C glucose feeding was chosen to both maximize the number of bacterial cells in the TEM sections and to reduce the dilution of ^13^C label due to metabolic turnover. For each timepoint and both cages (^13^C and ^12^C glucose treatment), the ileum of four bees were dissected. One gut was immediately preserved for electron microscopy and NanoSIMS, while the other three guts were homogenized and aliquoted for CFU plating and metabolite extraction **(Fig. 3)**. The control samples (from the cage provided with ^12^C glucose) were used to determine the natural ^13^C abundance in the bee tissue and *S. alvi* cells. Image metadata and Raw NanoSIMS region of interest (ROI) values can be found on github: https://github.com/Yelchazli/Foraging-on-host-synthesized-metabolites-enables-S.-alvi-to-colonize-the-honey-bee-gut.

### Quantification of bacterial loads via qPCR

DNA extraction was carried out by adding 250 μL of 2x CTAB to frozen gut homogenates. The mixture was homogenized at 6 m/second for 45 seconds in a Fast-Prep24TM5G homogenizer (MP Biomedicals) and centrifuged for 2 minutes at 2000 rpm. DNA was extracted in two-steps with Roti-Phenol and Phenol:Chloroform:Isoamylalcohol (25:24:1), then eluted in 200 μL DNAse/RNAse water and frozen. qPCR was run using target-specific primers for *Apis mellifera* actin, and specific 16S rRNA gene primers for *S. alvi* and *Gilliamella spp.* as well as 16S rRNA gene universal primers used in ^12^. qPCR reactions were run on a QuantStudio 3 (Applied Biosystems), and standard curves were performed using the respective amplicons on a plasmid as previously described ^12^.

In addition, 18S rRNA gene amplification with universal primers were run to assess fungal loads (F: TATGCCGACTAGGGATCGGG, R: CTGGACCTGGTGAGTTTCCC) ^42^. qPCR reactions were run on 384-well plate QuantStudio 5 (Applied Biosystems). The corresponding standard curve for absolute quantification was performed using the 199 bp amplicon as opposed to a plasmid containing the amplicon. Data was extracted and analyzed as previously described ^12^.

### Metabolite extractions

Frozen gut samples were divided and extracted via two established methods for short chain fatty acids (SCFAs) and soluble metabolites. Prior to soluble metabolite extraction from pure pollen grains, an equivalent amount to what was digested per bee was mixed with PBS. In order to approximate physio-chemical degradation of the pollen in the bee gut, the samples were bead beaten and incubated at 35°C for 18 hours.

SCFAs were extracted with 750 μL diethyl ether from 75 μL of gut supernatant that was first acidified with 5 μL of 11% HCl. Isovalerate [200 μM] (Sigma Aldrich) was used as an internal standard. The solvents and resulting two-phase mixture were kept cold at −20 °C and vortexed 3x for 30 seconds. The samples were then centrifuged for 3 minutes at 4 °C and 13000 g^-1^, and 80 μL of the ether phase was removed to a glass GC vial (Agilent). The sample was derivatized with 20 μL MTBSTFA + 1% TBDMS (Sigma Aldrich) for 1 hour at 30 °C ^43^.

Soluble metabolites were extracted via a modified Bligh and Dyer protocol ^44^. A (5:2:1) mixture of methanol (Sigma Aldrich) and chloroform (Sigma Aldrich) and double distilled water was chilled to −20 °C and 800 μL added to 100 μL of gut homogenate. A mixture of norleucine (Sigma Aldrich), norvaline (Sigma Aldrich), and ^13^C glucose (Cambridge Isotopes) was used as internal standards. The samples were vortexed three times for 30 seconds and extracted at −20 °C for 90 minutes. The samples were then centrifuged at 13,300 g^-1^ and 4 °C for 5 minutes. Liquid supernatant was transferred to a new tube and 400 μL of cold chloroform : methanol (1:1) was added to the insoluble material. The samples were again vortexed three times, extracted for 30 minutes at −20 °C, centrifuged, and the two supernatants were combined. Phase separation was achieved by adding 200 μL of water and centrifuging at the same conditions. The upper phase was then dried overnight in a speed vacuum. The sample was derivatized with 50 μL of 20 mg/mL methoxyamine hydrochloride in pyridine (Sigma Aldrich), for 90 minutes at 33 °C followed by silylation with 50 μL of MSTFA (Sigma Aldrich) for 120 minutes at 45 °C.

### GC-MS analysis

Samples were analyzed on an Agilent 8890/5977B series GC-MSD equipped with an autosampler that injected 1 μL of sample onto a VF-5MS (30m x 0.25 mm x 0.25 um) column. The SCFA samples were injected with a split ratio of 25:1, helium flow rate of 1 mL/min, and inlet temperature of 230 °C. The oven was held for 2 minutes at 50 °C, raised at 25 °C/min to 175 °C and then raised at 30 °C/minute to 280 °C and held for 3.5 minutes. The MSD was run in SIM mode with 3 target ions for each compound. The Soluble metabolite samples were injected with a split ratio of 15:1, helium flow rate of 1 mL/min and inlet temperature of 280 °C. The oven was held for 2 minutes at 125 °C, raised at 3 °C/minute to 150 °C, 5 °C/minute to 225 °C, and 15 °C/minute to 300 °C and held for 1.3 minutes. The MSD was run in scan mode from 50-500 Da at a frequency of 3.2 scan/second.

### Metabolomics data analysis

Mass spectrometry features were identified by spectral matching to the NIST17 mass spectra library or to in house analytical standards. Features were then extracted using the MassHunter Quantitative Analysis software (Agilent). Data normalization and analysis was performed using custom R scripts (See Data availability). Absolute metabolite quantification was performed by fitting normalized response values to calibration curves of analytical standards. The effect of diet, and colonization with *S. alvi* and *Gilliamella* were assessed by fitting z-score normalized metabolite abundances to a mixed linear model using the lmm2met package in R with diet, *S. alvi* and *Gilliamella* as fixed effect factors and experimental replicate as a random effect ^45^. The significance of each fixed effect was then adjusted by the Benjamini, Hochberg, and Yekutieli method to control for false discovery. Mass isotopologues were adjusted to account for natural isotope abundances following published procedures ^46,47^.

### TEM section preparation and NanoSIMS analysis

Bee guts were dissected, the ileum was cut away from the rest of the gut and fixed in glutaraldehyde solution (EMS, Hatfield, PA, US) 2.5% in Phosphate Buffer (PB 0.1M pH7.4) (Sigma, St Louis, MO, US) for 2 hours at room temperature (RT) and kept at 4°C until sample preparation. Ileum pieces were rinsed 3 times for 5 minutes in PB buffer and then postfixed with a fresh mixture of osmium tetroxide 1% (EMS, Hatfield, PA, US) with 1.5% potassium ferrocyanide (Sigma, St Louis, MO, US) in PB buffer for 2 hours at RT. The samples were then washed three times in distilled water and dehydrated in acetone solution (Sigma, St Louis, MO, US) at graded concentrations (30%-40min; 70%-40min; 100%-1h; 100%-2h). Dehydration was followed by infiltration in Epon resin (Sigma, St Louis, MO, US) at graded concentrations (Epon 1/3 acetone-2 hours; Epon 3/1 acetone-2 hours, Epon 1/1-4 hours; Epon 1/1-12 hours) and finally polymerized for 48 hours at 60°C in an oven. Thin sections of 100 nanometers (nm) were cut on a Leica Ultracut (Leica Mikrosysteme GmbH, Vienna, Austria) and picked up on copper slot grid 2×1mm (EMS, Hatfield, PA, US) coated with a Polyetherimide (PEI) film (Sigma, St Louis, MO, US). Sections were poststained with uranyl acetate (Sigma, St Louis, MO, US) 2% in H_2_O for 10 minutes, rinsed several times with H_2_O followed by Reynolds lead citrate for 10 minutes and rinsed several times with H_2_O.

Large montages with a pixel size of 6.9 nm covering areas of around 120×120 µm were taken with a transmission electron microscope Philips CM100 (Thermo Fisher Scientific, Hillsboro, USA) at an acceleration voltage of 80kV with a TVIPS TemCam-F416 digital camera (TVIPS GmbH, Gauting, Germany) and the alignment was performed using Blendmont command-line program from the IMOD software ^48^.

Four holes were made using the electron beam at the four corners of the montage to localize the area of interest for NanoSIMS imaging. Then the slot grid was deposited on 1 µL of distilled water on a 10 mm round coverslip and left to dry for 10 minutes. The slot grid was detached, letting the section attach to the coverslip.

After being coated with a 15nm gold layer to evacuate a potential build-up of a charge during the NanoSIMS analyses, the samples were analyzed by a NanoSIMS 50L (CAMECA). A primary beam of 16 keV Cs+ ions was focused to a spot size of about 100 nm on the sample surface, causing the sputtering of atoms and small molecules from the first 5-10 nm of the sample. Negative secondary ions were directed and focalized into a single beam before reaching the magnetic sector of the mass spectrometer where the ions were separated. The species of interest (^12^C^14^N-and ^13^C^14^N-for this study) were then measured simultaneously using Electron Multipliers. To resolve the interference between ^13^C^14^N-and ^12^C^15^N-, two slits (ES and AS) were used to achieve a mass resolution higher than 10000 (Cameca definition), ^49^.

In the imaging mode, selected areas of 25×25 µm were implanted using a strong beam (using diaphragm D1-1 (diameter 750 µm) and a current of 16 pA) to remove the gold coating and reach stable emission conditions. After optimization of the extraction of the secondary ions, the area was imaged by scanning with a smaller beam (∼100 nm, using diaphragm D1-5 (diameter 100 µm) and a current of 0.45 pA) with a 256×256 pixel resolution and a dwell time of 5µs/pixel. For each image, 10 layers were acquired. The image processing was performed using “L’image” software (Larry Nittler, Carnegie Institution of Washington). The 10 layers were aligned and stacked, 44 ms deadtime correction was applied. Regions of interests (ROI) were manually drawn around bacterial cells, gut epithelium layer, and bulk host cells. The resulting data is reported in **Dataset S1**. At% enrichment was calculated in R using the data.table package. At% enrichment formula: 13*C At*% = ^14N 13C^/_(14N 13C + 14N 12C)_ × 100. **(Fig. S9).**

### *In vitro* growth of *S. alvi*

*S. alvi* was cultured in Bee9 media, an M9-derived minimal defined media containing essential vitamins and mixtures of organic acids (see recipe in **Dataset S3** on github: https://github.com/Yelchazli/Foraging-on-host-synthesized-metabolites-enables-S.-alvi-to-colonize-the-honey-bee-gut). Growth assays were performed in 96-well plates (Corning costar) at 35 °C, 5% CO_2_, with continuous orbital shaking (H1 Synergy, Biotek). OD_600_ measurements were taken every 30 minutes for 20 hours. All conditions were independently replicated three times. For metabolomics measurements, plates were immediately placed on ice at the end of each experiment, and liquid was transferred to microcentrifuge tubes. The tubes were then centrifuged at 13,300 g and 4 °C for 30 seconds, and the liquid supernatant was then snap frozen in liquid nitrogen and stored at −80 °C for metabolomic analysis.

### Phylogenetic analysis of the kynureninase gene family

The kynureninase protein sequence of *S. alvi* wkB2 was retrieved from NCBI (WP 025330329.1), and blasted (NCBI, BlastP) against the “nr” database. Based on the BlastP hits, 227 divergent amino acid sequences were exported in fasta format and aligned using MUSCLE ^50^ with standard parameters. Tree inference and bootstrap validation were performed using IQtree with the following command: “<u>iqtree-</u> omp -s kyn_sequences_muscle.aln -nt 6 -m TEST -bb 1000” ^51^. The tree topology was visualized using the ITOL online viewer (https://itol.embl.de/).

## Data Availability

Raw data and scripts used to process data and generate figures are available on github at the following link: https://github.com/Yelchazli/Foraging-on-host-synthesized-metabolites-enables-S.-alvi-to-colonize-the-honey-bee-gut. Metabolomics data have been deposited to the EMBL-EBI MetaboLights database (DOI: 10.1093/nar/gkz1019, PMID:31691833) with the identifier MTBLS8058. https://www.ebi.ac.uk/metabolights/MTBLS8058

## Acknowledgments

We would like to thank Silvia Moriano-Gutierrez, and Gonçalo Santos-Matos for providing helpful comments on the manuscript. This work was funded by the University of Lausanne, an ERC Starting Grant (MicroBeeOme), the NCCR Microbiomes, a National Centre of Competence in Research, funded by the Swiss National Science Foundation (grant no. 180575) and a Swiss National Science Foundation project grant (grant no. 179487) to P.E.

## Author Contributions

A.Q., Y.E. and P.E. conceived the project and designed the experiments. A.Q, and Y.E. performed the experiments. A.Q. analyzed the GC-MS results, while Y.E. analyzed the qPCR and NanoSIMS results. N.N. assisted with bee experiments, metabolite, and DNA extractions. A.M.N. performed preliminary in vitro growth experiments. J.D. performed the tissue sectioning and TEM imaging supervised by C.G. S.E. performed the NanoSIMS imaging and instructed on the data analysis, supervised by A.M., who also advised on the design of the NanoSIMS experiment. A.Q., Y.E. and P.E. wrote the manuscript, with contributions from S.E. and A.M. All authors reviewed the manuscript and provided feedback. As equally contributing first authors, A.Q. and Y.E. may each list their name first when referencing this work.

## Supplementary Figures and Tables

**Figure S1.**
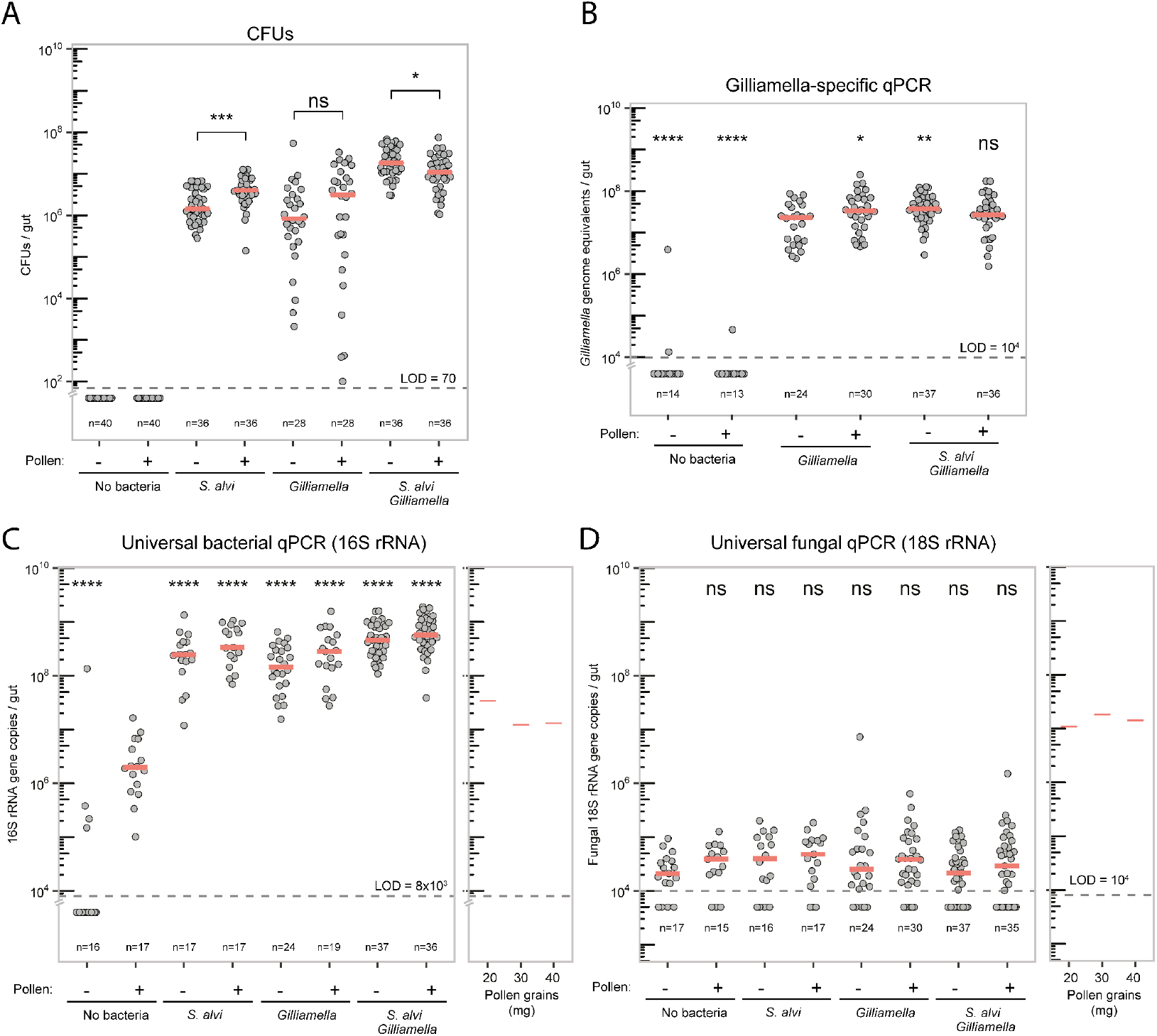
Total number of bacteria and fungi and colonization levels of *Gilliamella* per bee gut. **(A)** Total number of bacteria (both *S. alvi* and *Gilliamella*) was determined by plating homogenized guts on BHIA and counting CFUs. Each dot represents a single bee gut and red bars represent the median value for each group. Horizontal dotted line represents LOD = < 70. No colonies were observed in samples plotted below the LOD line. Wilcoxon rank sum test was done with *S. alvi* (Pollen -) as reference group, ns, nonsignificant; ****P < 0.001. (**B**) Bacterial cells (number of bacterial genome equivalents) per bee gut determined by qPCR using genus-specific *Gilliamella* 16S rRNA gene primers. Each dot represents the number of cells in a single bee gut, red bars represent the median, horizontal black line represents qPCR LOD =< 10^4^. The presence of pollen in the diet is indicated by (P). Wilcoxon rank sum test was done with *Gilliamella* (Pollen -) as reference group, ns, nonsignificant; ****P < 0.001. (**C**) Number of 16S rRNA gene copies per bee gut and in three different quantities of sterilized bee pollen as determined by qPCR using universal 16S rRNA gene primers. Each dot represents the 16S rRNA gene copies in a single bee gut, red bars represent the median, horizontal black line represents qPCR LOD = < 8×10^3^. The presence of pollen in the diet is indicated by (P). Wilcoxon rank sum test was done with MF (Pollen -) as reference group, ns, nonsignificant; ****P < 0.001. The number of 16S rRNA gene copies detected in DNA extracted from different quantities of sterile pollen used in the bee diet is shown in a separate plot. The high background signal in samples containing pollen comes from the amplification of chloroplast DNA as shown in previous studies ^52^. (**D**) Number of 18S rRNA gene copies per bee gut and in three different quantities of sterilized bee pollen as determined by qPCR using universal 18S rRNA gene primers. Each dot represents the 18S rRNA gene copies in a single bee gut, red bars represent the median, horizontal black line represents qPCR LOD = < 10^4^. The presence of pollen in the diet is indicated by (P). Wilcoxon rank sum test was done with MF (Pollen -) as reference group, ns, nonsignificant; ****P < 0.001.

**Figure S2.**
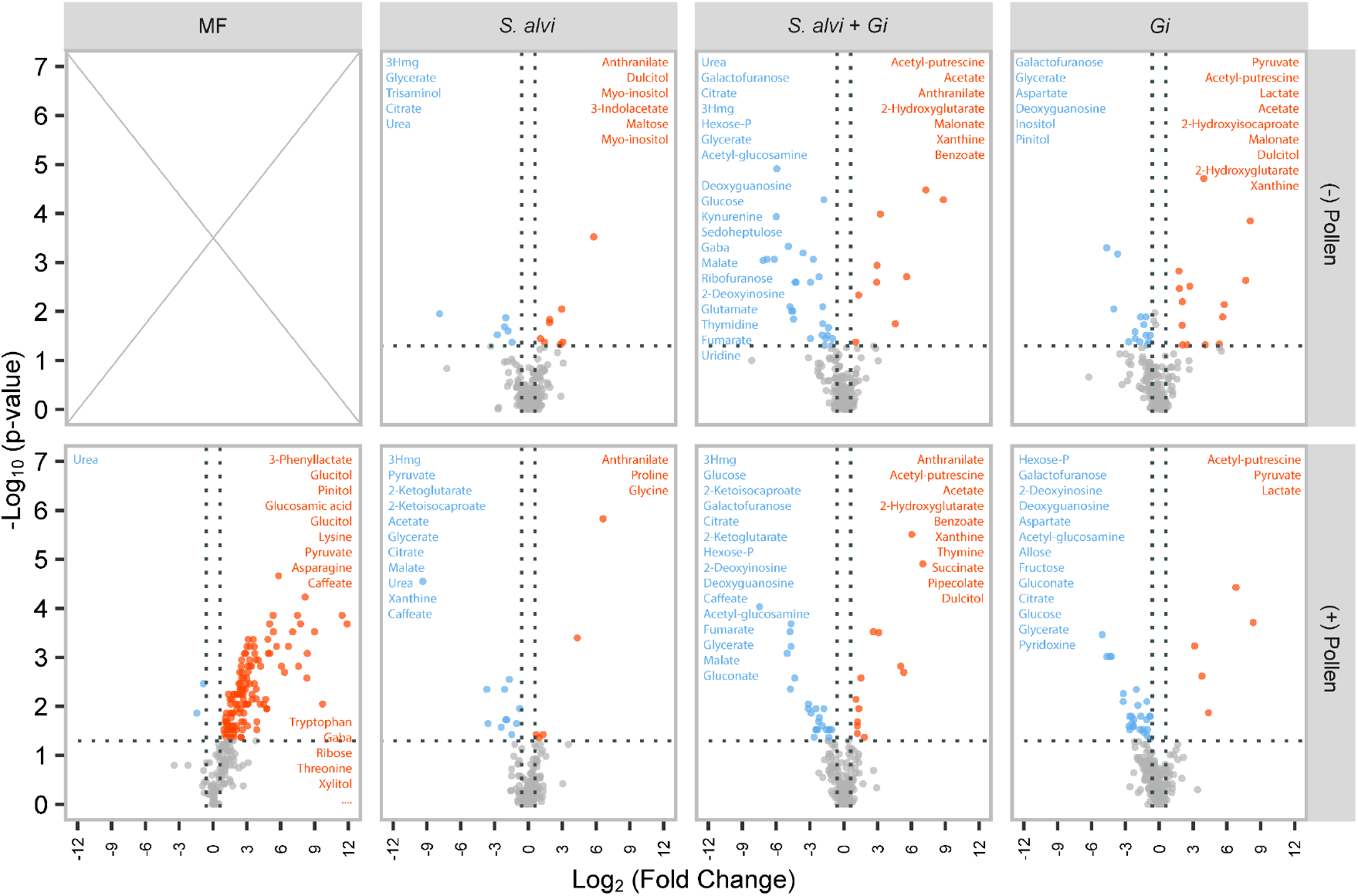
Volcano plots highlighting gut metabolite abundance changes between colonized and microbiota-free (MF) bees with and without pollen. Significantly differentially abundant metabolites between mono-and co-colonized *S. alvi* and *Gilliamella (Gi)* vs MF bees, as well as between MF ((+) Pollen) and MF ((-) Pollen) bees. Significantly depleted and produced metabolites are shown in blue and orange, respectively. Annotations are listed in order of significance for identified metabolites, with isomers and other duplications removed. The full set of significantly more abundant metabolites in the first panel can be found in the supporting scripts. Adjusted significance values were calculated using Wilcoxon rank sum test, and Benjamini-Hochberg correction, using the MF conditions as a reference.

**Figure S3.**
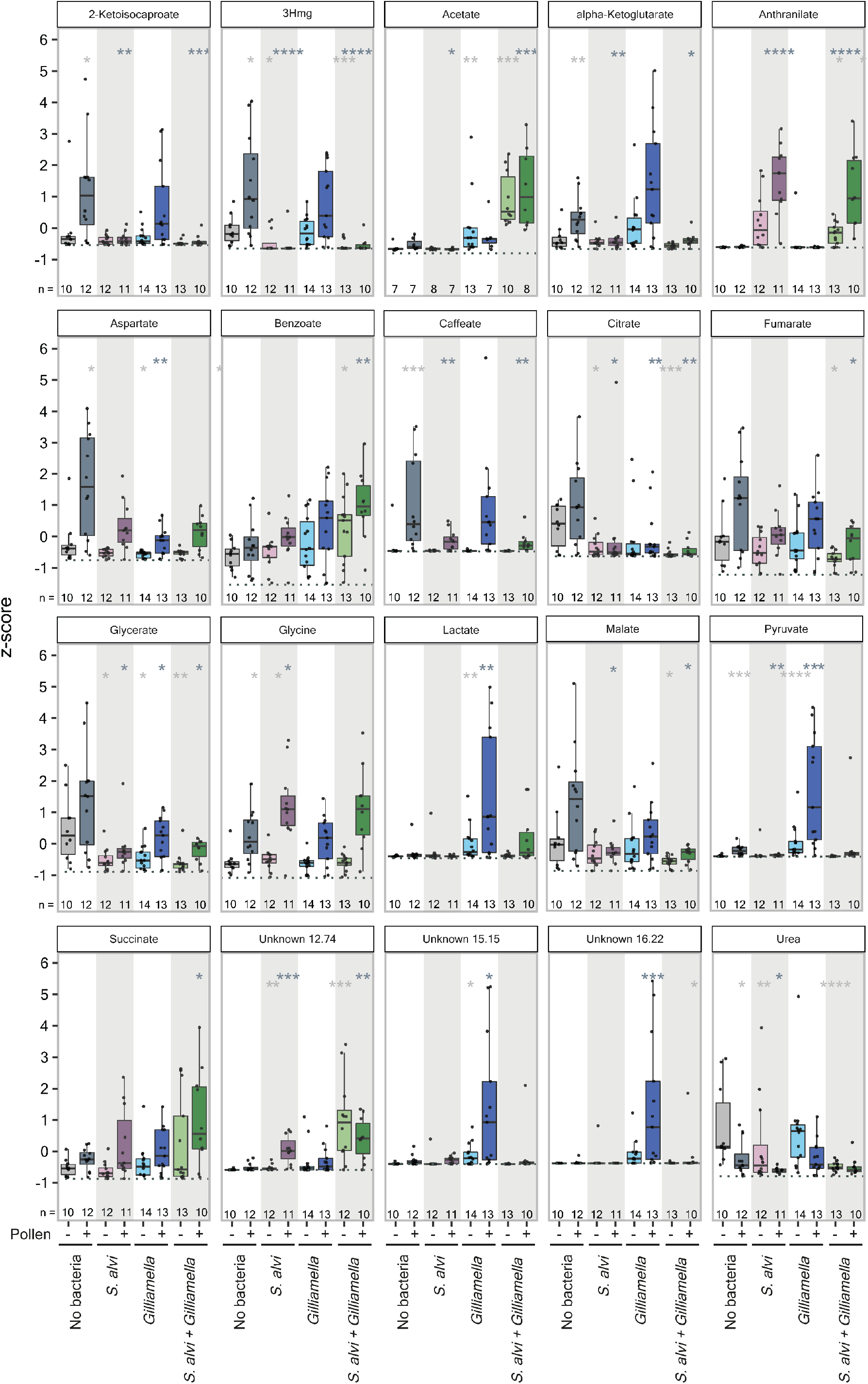
Z-score normalized metabolite abundances of compounds significantly affected by *S. alvi* colonization. Box plots and individual data points display values across all 8 experimental conditions. Only metabolites in which a mixed linear model showed a significant effect of *S*. *alvi* colonization are plotted. Multiple types of metabolic interactions are shown. For example, metabolite depletion by *S. alvi* exclusively (2-ketoisocaproate, 3Hmg, caffeate, urea) or by both species (aspartate, citrate, glycerate), metabolite production by *S. alvi* exclusively (anthranilate, unknown 12.74), cross-feeding from *Gilliamella* to *S. alvi* (pyruvate, lactate), synergistic metabolite depletion (fumarate, malate) and production (acetate, benzoate). The light and dark gray asterisks show groups significantly different from MF bees fed sugar water and pollen or only sugar water (also indicated by +/-sign), calculated using Wilcoxon rank sum test, * P < 0.05, ** < 0.01, *** < 0.001, **** < 0.0001. The dotted line represents the transformed LOD for each metabolite.

**Figure S4.**
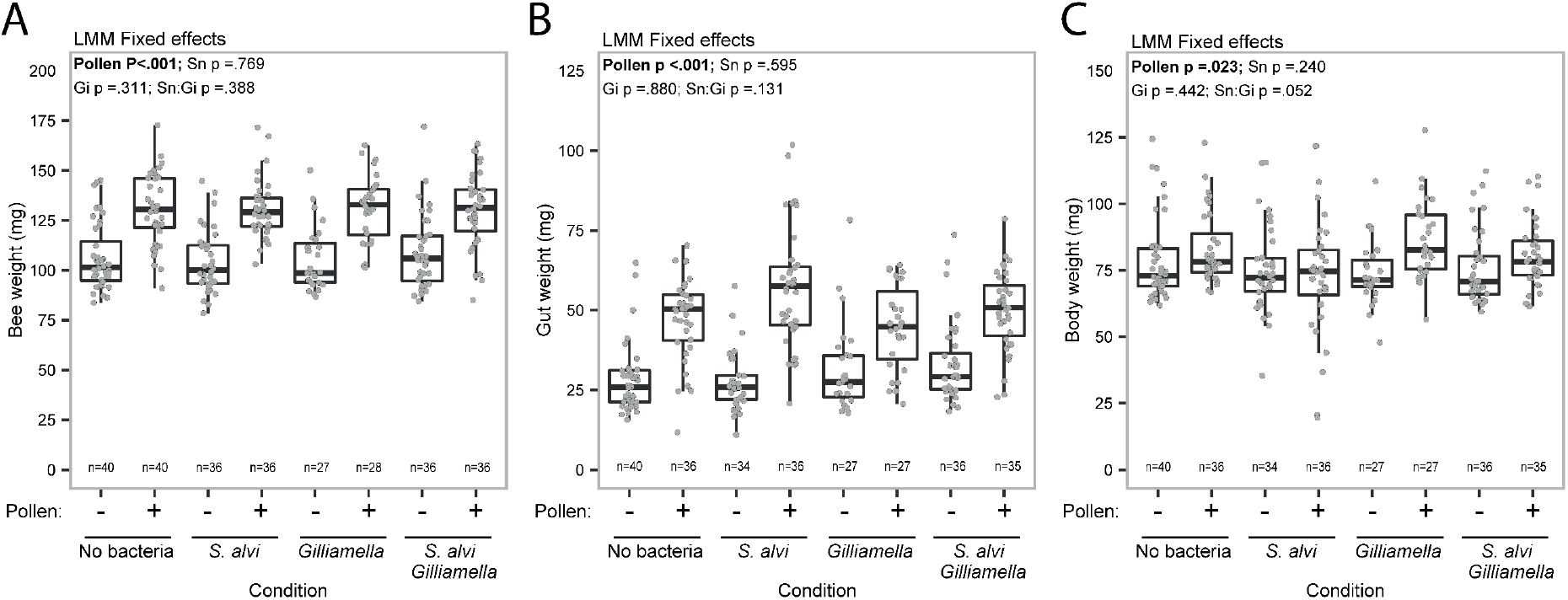
Pollen consumption but not colonization increases bee gut weight. **(A) Whole bee and** (B) dissected bee gut weights are strongly influenced by pollen in the diet, but not by colonization treatment. (**C**) The body weight, or difference between bee and gut weights, was also slightly higher across colonization groups for bees fed a pollen diet. The p-values (ANOVA) are shown for the fixed effect terms in the linear mixed models fitted by ML with nested, random cage effects.

**Figure S5.**
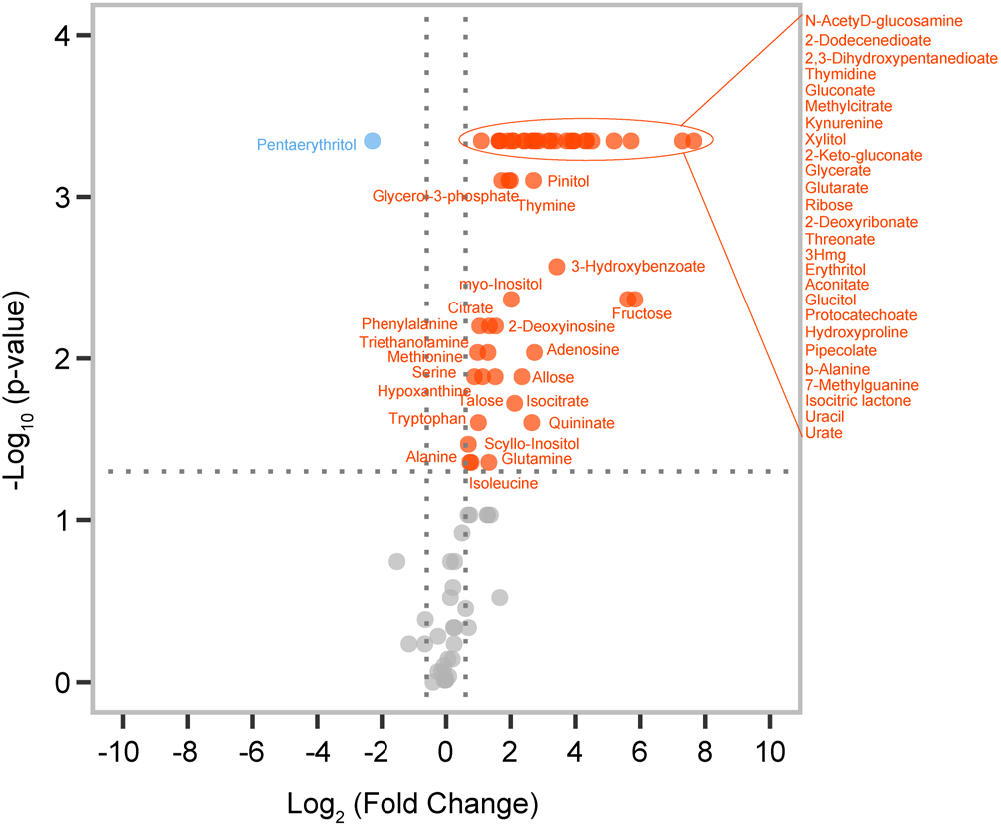
Volcano plot highlighting gut metabolic changes between Day 0 and Day 6 after emergence in MF bees without pollen. Many metabolites are more abundant in the gut on Day 6 than on Day 0 after emergence. Significantly less and more abundant metabolites on Day 6 are shown and annotated in blue and orange, respectively, with isomers and other duplications removed. Adjusted significance values were calculated using Wilcoxon rank sum test, and Benjamini-Hochberg correction, using the day zero conditions as a reference (n=8).

**Figure S6.**
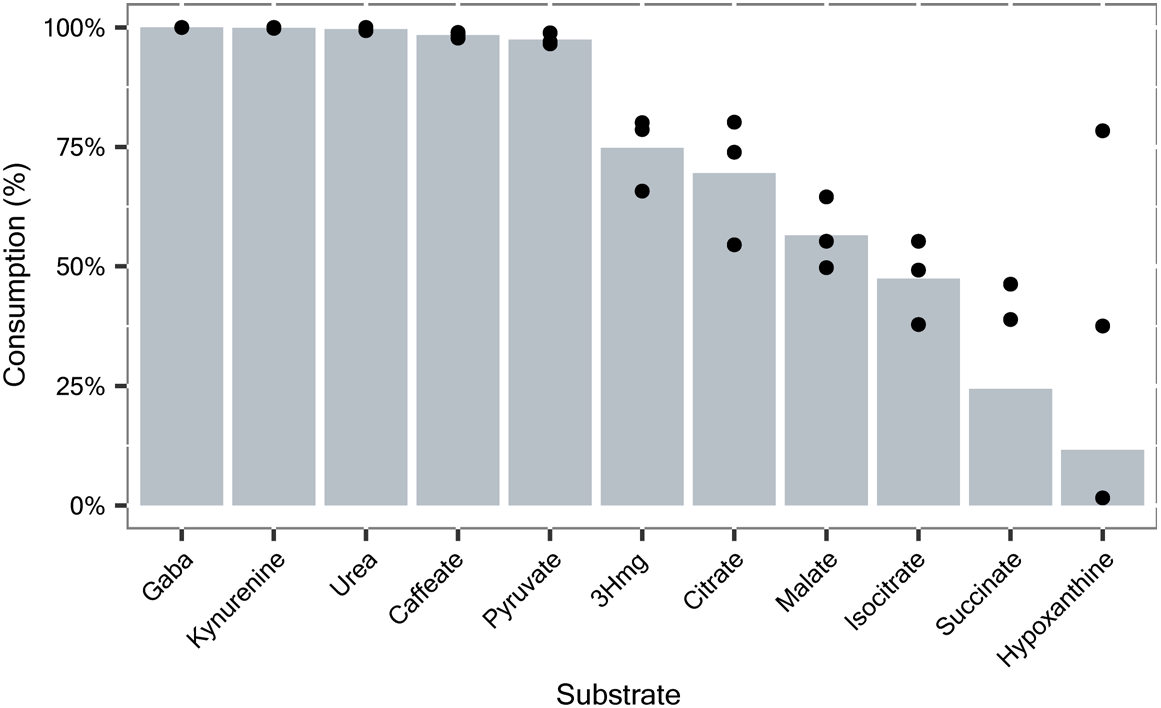
*In vitro* consumption of carbon substrates. The percentage of each carbon source consumed by the end of the *in vitro* growth experiment is shown for each tested carbon source. Data was taken from the condition in which all carbon sources were added to Bee9 media in equimolar amounts. Bars represent the average value from 3 biological replicates (black dots).

**Figure S7.**
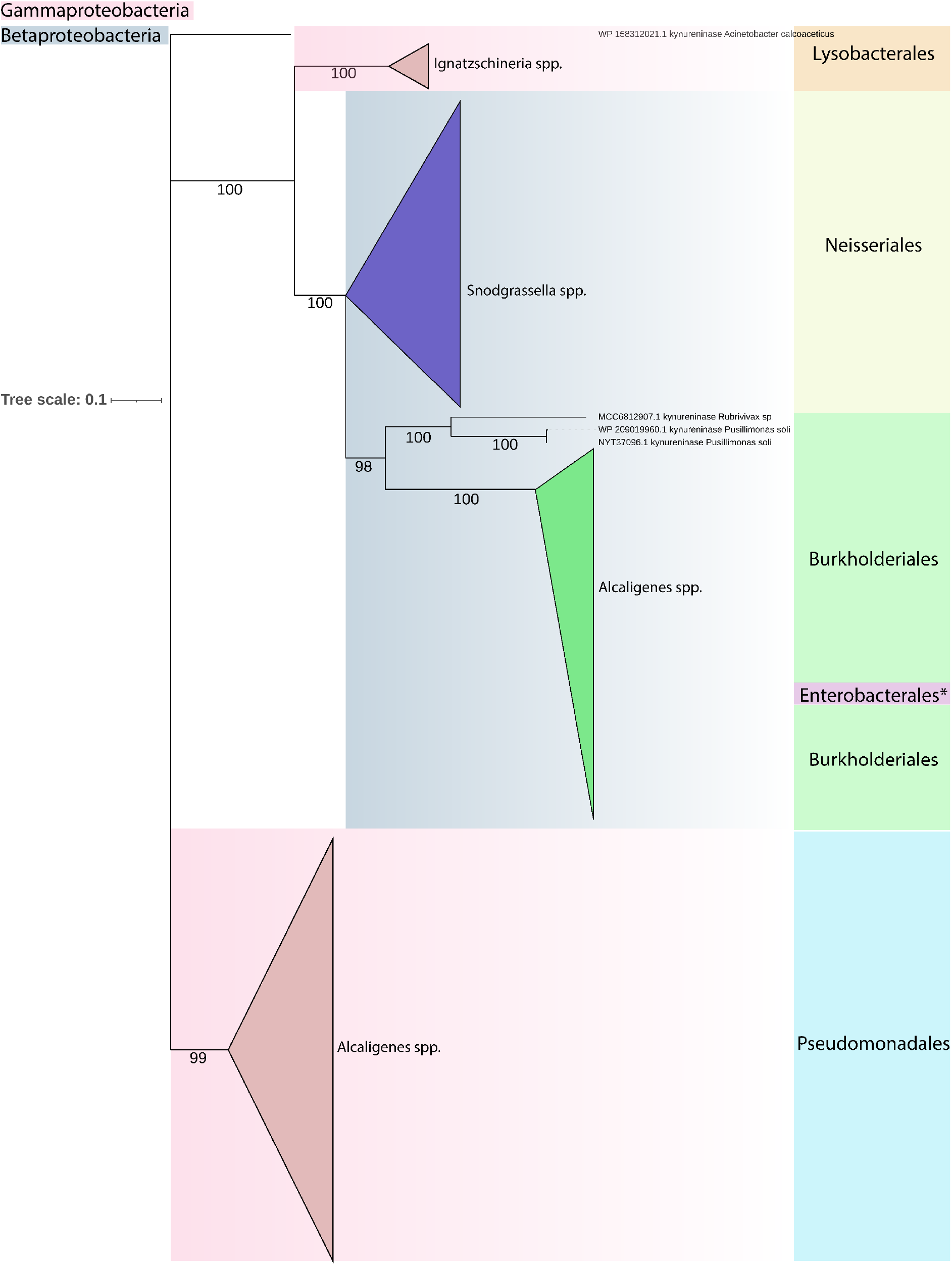
Phylogenetic tree of the kynureninase gene family found in *S. alvi*. The amino acid sequences of 227 homologs of the kynureninase gene identified in *S. alvi* wkb2 were retrieved from NCBI. The tree was inferred with Iqtree after aligning the sequences with MUSCLE (see methods). Tips are collapsed for clades that contain sequences of the same genus. Colored boxes on the right indicate the bacterial family. Bootstrap values are indicated before every node. Gammaproteobacteria are highlighted in pink and Betaproteobacteria in blue. Asterisk indicates a sequence (MBY6345856.1) from Providencia rettgeri, a Gammaproteobacterium, which clusters together with homologs of the kynureninase of Betaproteopbacteria.

**Figure S8.**
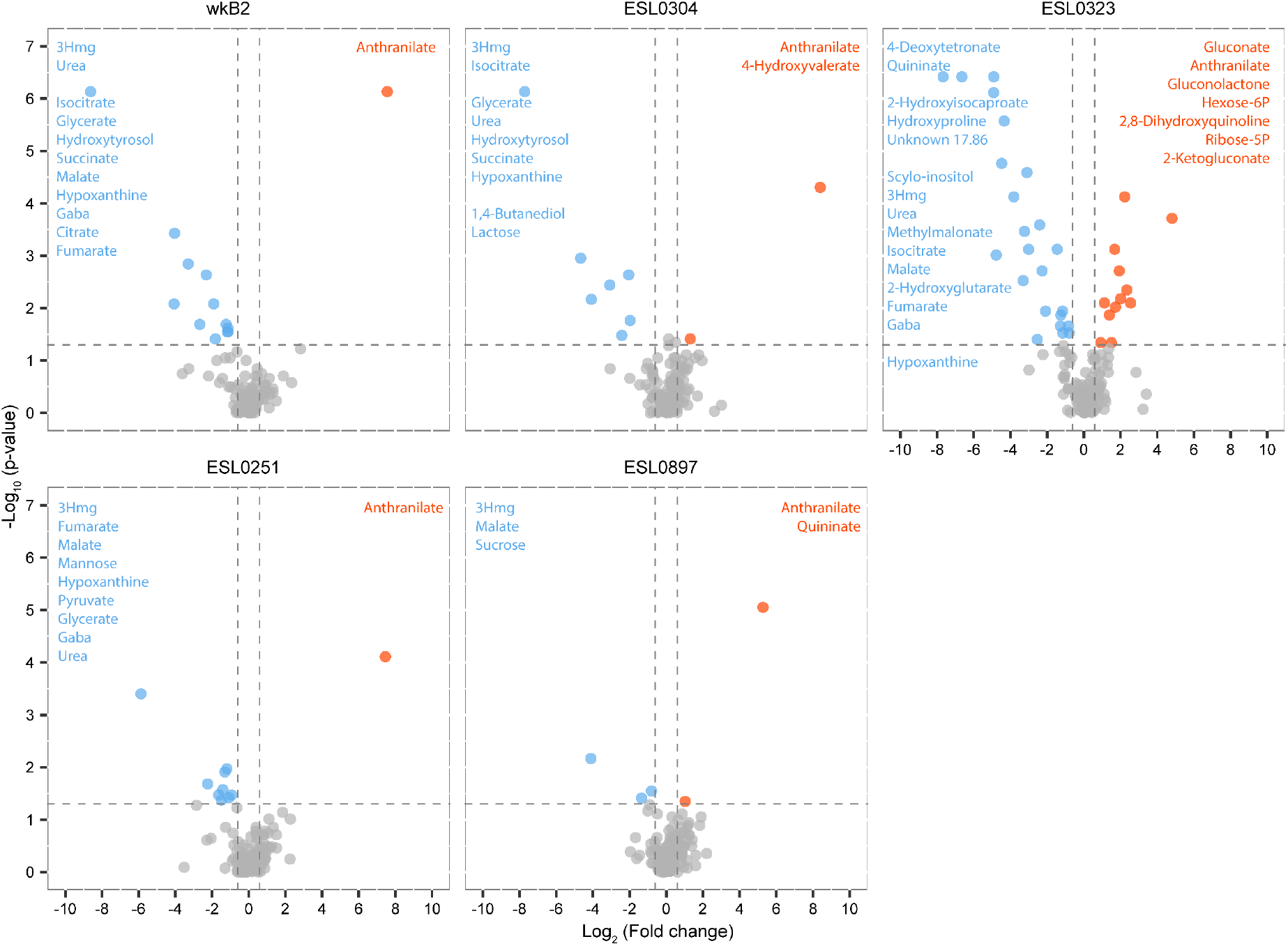
Gut metabolite changes in bees mono-colonized with divergent *Snodgrassella* strains relative to microbiota-free (MF) bees. Volcano plot of gut metabolites showing the similarities in metabolite changes from colonized vs MF bees across strains. Top: Strains native to *A. mellifera*; Bottom: Strains native to *Bombus sp.* Significantly depleted and produced metabolites are shown in blue and orange, respectively. Adjusted significance values were calculated using Wilcoxon rank sum test, and Benjamini-Hochberg correction, using the MF condition as a reference.

**Figure S9.**
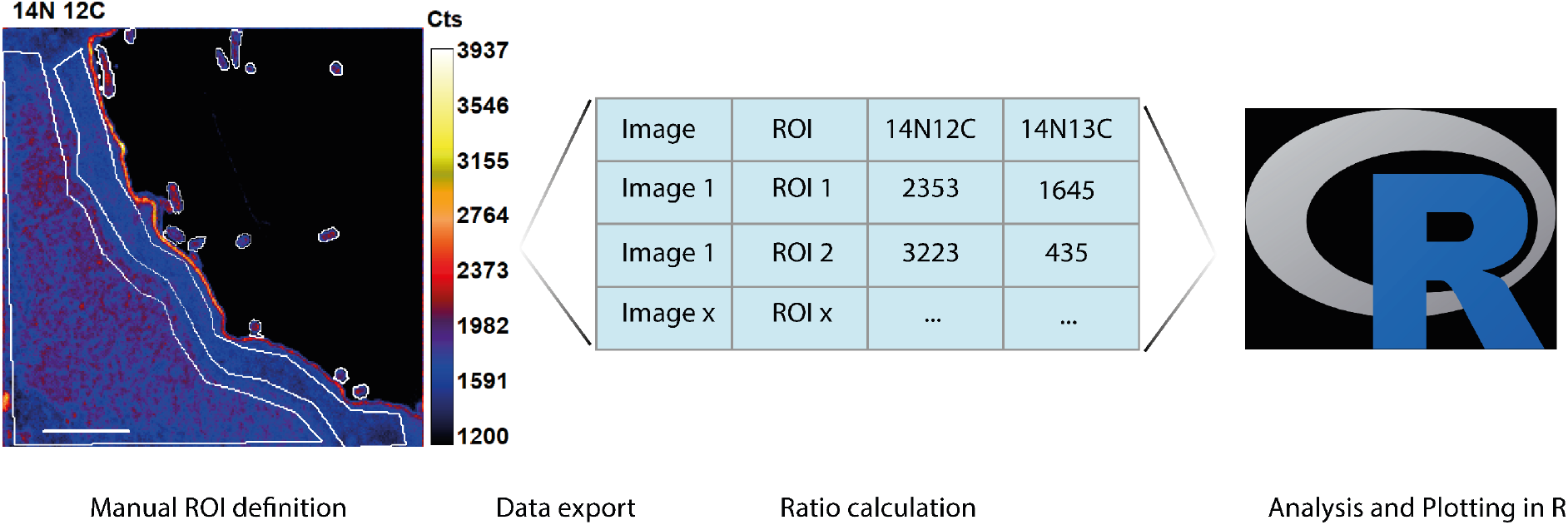
NanoSIMS image analysis workflow. For each image, 10 layers were acquired by bombarding the surface of the samples with Cs+ Ion beam (See methods). The image processing was performed using “L’image” software (Larry Nittler, Carnegie Institution of Washington). The 10 layers were aligned and stacked and 44 ms deadtime correction was applied. Regions of interests (ROI) were manually drawn around bacterial cells, gut epithelium layer, and bulk host cells, using a minimum threshold specified and values were extracted and reported in **Dataset S1**. At% enrichment was calculated in R using the data.table package. At% enrichment formula: 13*C At*% = ^14N 13C/^(14N 13C + 14N 12C) ^× 100^.

**Table S1.** Bacterial strains used in this study. ^12,53,54^.

| **Bacterial strain** | **# 16S rRNA copies** | **Culturing condition** | **Strain source** | **Place of origin** |
| --- | --- | --- | --- | --- |
| *Gilliamella apicola* wkB1<sup>T</sup> | 4 | BHIA, 35°C, microaerophilic | (Kwong & Moran, 2013) | New Haven, USA |
| *Gilliamella apis* ESL0169 | 4 | BHIA, 35°C, microaerophilic | (Kešnerová et al., 2017) | Lausanne, Switzerland |
| *Gilliamella sp.* ESL0182 | 4 | BHIA, 35°C, microaerophilic | (Ellegaard & Engel, 2018) | Lausanne, Switzerland |
| *Gilliamella apis* ESL0297 | 4 | BHIA, 35°C, microaerophilic | This study | Lausanne, Switzerland |
| *Snodgrassella alvi* wkB2<sup>T</sup> | 4 | TSA, 35°C, microaerophilic | (Kwong & Moran, 2013) | New Haven, USA |
| *Snodgrassella alvi* ESL0304 | 4 | TSA, 35°C, microaerophilic | This study | Lausanne, Switzerland |
| *Snodgrassella alvi* ESL0323 | 4 | TSA, 35°C, microaerophilic | This study | Lausanne, Switzerland |
| *Snodgrassella alvi* ESL0251 | 4 | TSA, 35°C, microaerophilic | This study | Lausanne, Switzerland |
| *Snodgrassella alvi* ESL0897 | 4 | TSA, 35°C, microaerophilic | This study | Lausanne, Switzerland |

**Table S2.** ***S. alvi* & *Gilliamella* colonization summary counts.** Total bee numbers per experiment, survival rate, number of bees used for CFU plating, qPCR and metabolomics analysis, and pollen consumption are shown.

|  | Replicate | 1 | 2 | 3 | 4 | 5 | Total |
| --- | --- | --- | --- | --- | --- | --- | --- |
| **Bees** | **MD** | 34 | 24 | 24 | 24 | 0 | 106 |
|  | **MD (P)** | 30 | 23 | 23 | 25 | 0 | 101 |
|  | **Sn** | 31 | 24 | 20 | 0 | 20 | 95 |
|  | **Sn (P)** | 34 | 23 | 20 | 0 | 19 | 96 |
|  | **Sn+Gi** | 32 | 22 | 24 | 0 | 16 | 94 |
|  | **Sn+Gi (P)** | 29 | 24 | 20 | 0 | 22 | 95 |
|  | **Gi** | 0 | 0 | 0 | 26 | 25 | 51 |
|  | **Gi (P)** | 0 | 0 | 0 | 25 | 25 | 50 |
| **Survival (%)** | **MD** | 100% | 100% | 100% | 96% | N/A | 99% |
|  | **MD (P)** | 100% | 100% | 100% | 100% | N/A | 100% |
|  | **Sn** | 94% | 100% | 100% | N/A | 100% | 98% |
|  | **Sn (P)** | 100% | 100% | 95% | N/A | 100% | 99% |
|  | **Sn+Gi** | 97% | 95% | 100% | N/A | 100% | 98% |
|  | **Sn+Gi (P)** | 97% | 92% | 95% | N/A | 100% | 96% |
|  | **Gi** | N/A | N/A | N/A | 100% | 100% | 100% |
|  | **Gi (P)** | N/A | N/A | N/A | 96% | 100% | 98% |
| **CFU** | **MD** | 10 | 10 | 10 | 10 | N/A | 40 |
|  | **MD (P)** | 10 | 10 | 10 | 10 | N/A | 40 |
|  | **Sn** | 10 | 10 | 10 | N/A | 6 | 36 |
|  | **Sn (P)** | 10 | 10 | 10 | N/A | 6 | 36 |
|  | **Sn+Gi** | 10 | 10 | 10 | N/A | 6 | 36 |
|  | **Sn+Gi (P)** | 10 | 10 | 10 | N/A | 6 | 36 |
|  | **Gi** | N/A | N/A | N/A | 10 | 18 | 28 |
|  | **Gi (P)** | N/A | N/A | N/A | 10 | 18 | 28 |
| **qPCR** | **MD** | 4 | 3 | 4 | 6 | N/A | 17 |
|  | **MD (P)** | 4 | 3 | 4 | 5 | N/A | 16 |
|  | **Sn** | 4 | 9 | 6 | N/A | 6 | 25 |
|  | **Sn (P)** | 4 | 9 | 8 | N/A | 6 | 27 |
|  | **Sn/Gi** | 10 | 10 | 11 | N/A | 6 | 37 |
|  | **Sn/Gi (P)** | 10 | 10 | 10 | N/A | 6 | 36 |
|  | **Gi** | N/A | N/A | N/A | 10 | 14 | 24 |
|  | **Gi (P)** | N/A | N/A | N/A | 9 | 21 | 30 |
| **Metabolomics** | **MD** | 2 | 1 | 2 | 5 | N/A | 10 |
|  | **MD (P)** | 2 | 1 | 4 | 5 | N/A | 12 |
|  | **Sn** | 1 | 1 | 2 | N/A | 6 | 10 |
|  | **Sn (P)** | 1 | 1 | 3 | N/A | 6 | 11 |
|  | **Sn+Gi** | 3 | 1 | 5 | N/A | 6 | 15 |
|  | **Sn+Gi (P)** | 1 | 1 | 4 | N/A | 5 | 11 |
|  | **Gi** | N/A | N/A | N/A | 5 | 9 | 14 |
|  | **Gi (P)** | N/A | N/A | N/A | 6 | 7 | 13 |
| **Pollen Consumption (mg/bee)** | **MD** | - | - | - | - | - | - |
|  | **MD (P)** | 30 | 33 | 32 | 13 | N/A | 26 |
|  | **Sn** | - | - | - | - | - | - |
|  | **Sn (P)** | 20 | 31 | 30 | N/A | 22 | 26 |
|  | **Sn+Gi** | - | - | - | - | - | - |
|  | **Sn+Gi (P)** | 19 | 30 | 33 | N/A | 16 | 24 |
|  | **Gi** | - | - | - | - | - | - |
|  | **Gi (P)** | N/A | N/A | N/A | 17 | 18 | 17 |

**Table S3.** Summary of the number of bees sampled per replicate and treatment for the mono-colonization experiment with divergent *Snodgrassella* strains. Total number of bees per experiment, survival rate, and number of bees used for CFU plating and metabolomics analysis are shown.

|  | Replicate | 1 | 2 | Total |
| --- | --- | --- | --- | --- |
| **Bees** | **MD** | 14 | 15 | 29 |
|  | **wkB2** | 14 | 14 | 28 |
|  | **251** | 16 | 15 | 31 |
|  | **304** | 14 | 15 | 29 |
|  | **323** | 14 | 15 | 29 |
|  | **892** | 14 | 15 | 29 |
|  | **897** | 15 | 14 | 29 |
| **Survival (%)** | **MD** | 93% | 100% | 96% |
|  | **wkB2** | 100% | 100% | 100% |
|  | **251** | 88% | 100% | 94% |
|  | **304** | 100% | 80% | 90% |
|  | **323** | 100% | 100% | 100% |
|  | **892** | 86% | 100% | 93% |
|  | **897** | 80% | 79% | 79% |
| **CFU** | **MD** | 12 | 12 | 24 |
|  | **wkB2** | 12 | 12 | 24 |
|  | **251** | 11 | 12 | 23 |
|  | **304** | 12 | 9 | 21 |
|  | **323** | 12 | 12 | 24 |
|  | **892** | 12 | 12 | 24 |
|  | **897** | 12 | 9 | 21 |
| **Metabolomics** | **MD** | 6 | 6 | 12 |
|  | **wkB2** | 6 | 6 | 12 |
|  | **251** | 8 | 9 | 17 |
|  | **304** | 6 | 6 | 12 |
|  | **323** | 7 | 6 | 13 |
|  | **892** | 6 | 6 | 12 |
|  | **897** | 6 | 6 | 12 |

