## Supplementary figures and images for "Foraging on host synthesized metabolites enables the bacterial symbiont *Snodgrassella alvi* to colonize the honey bee gut"

### Supplementary Dataset 2

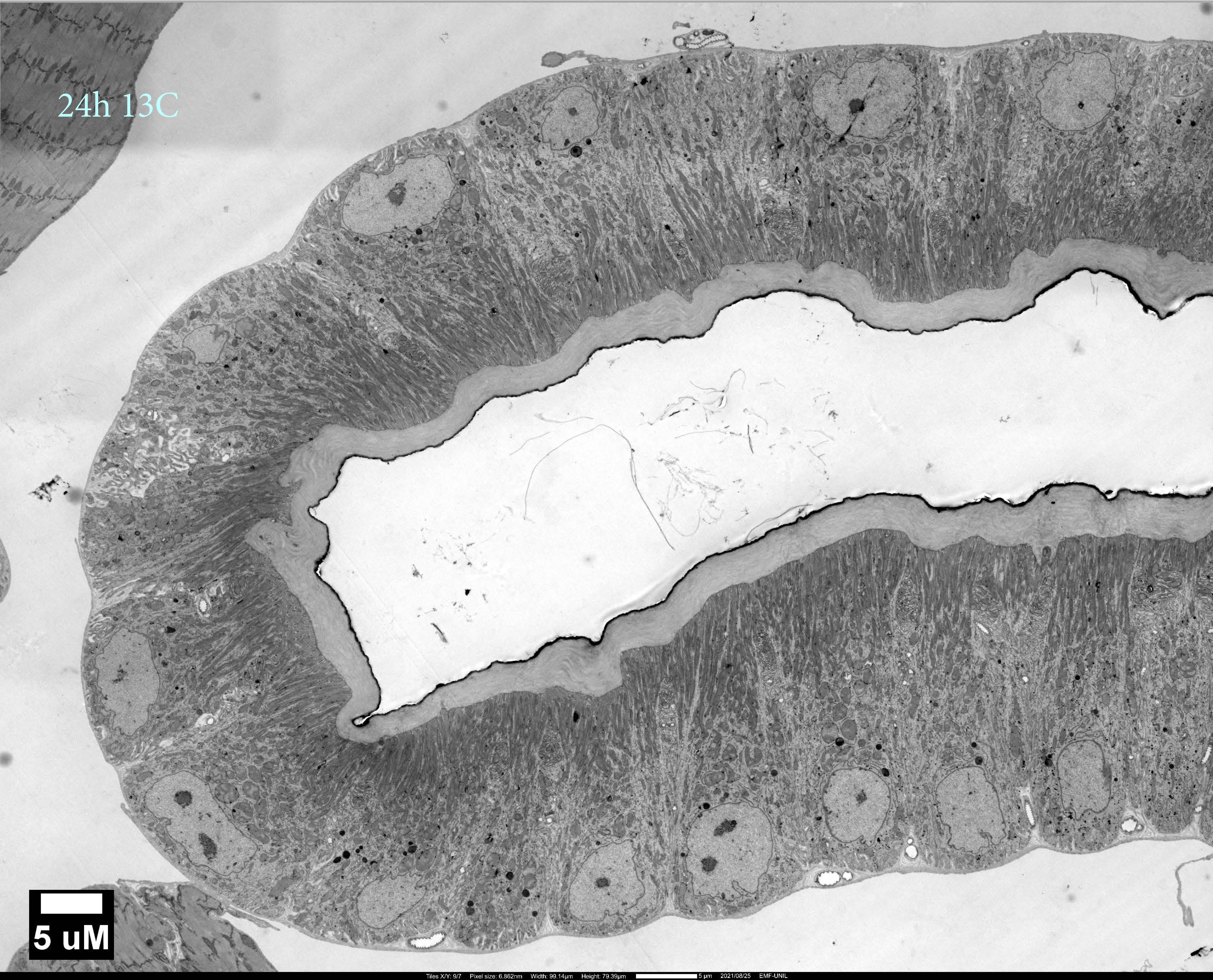

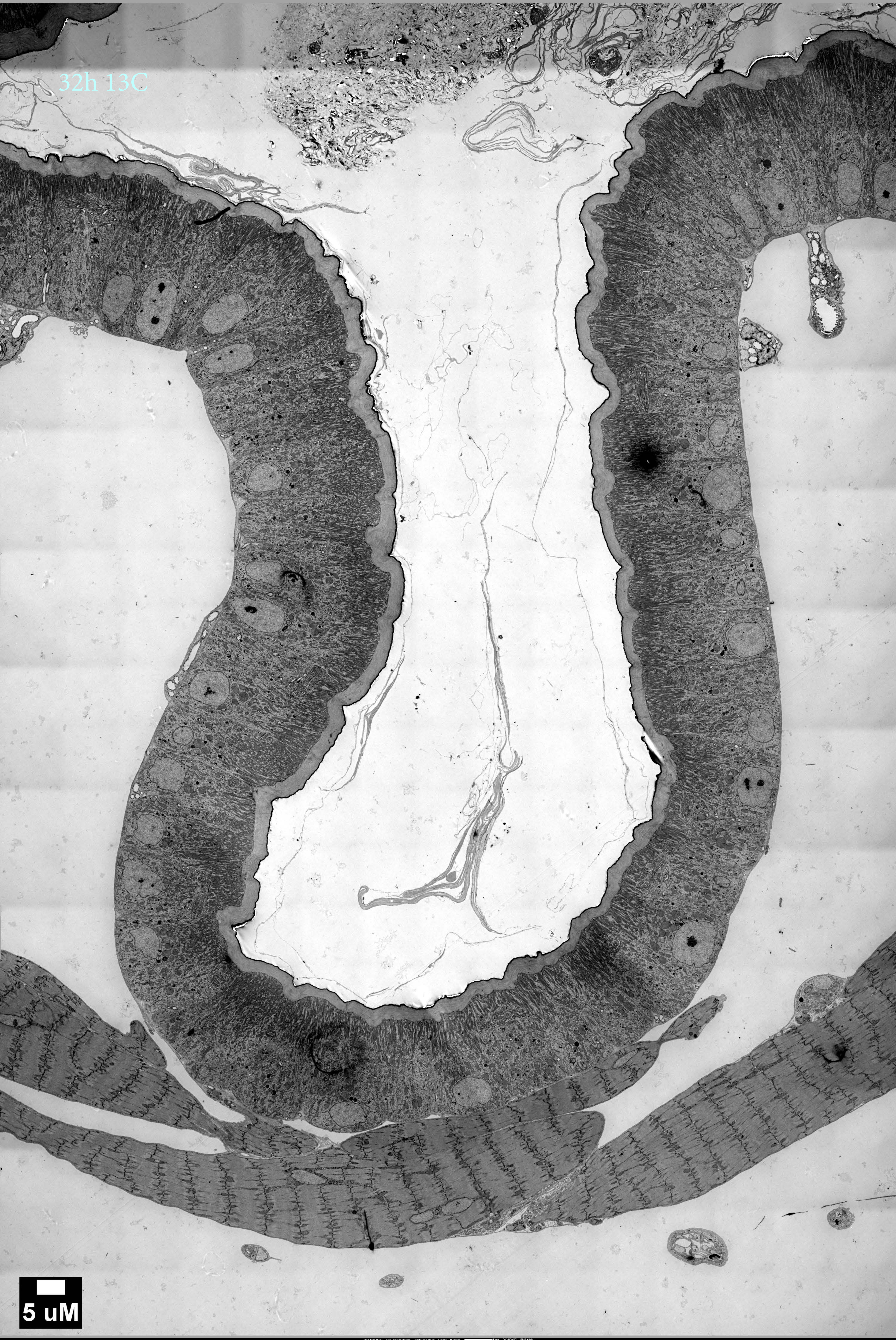

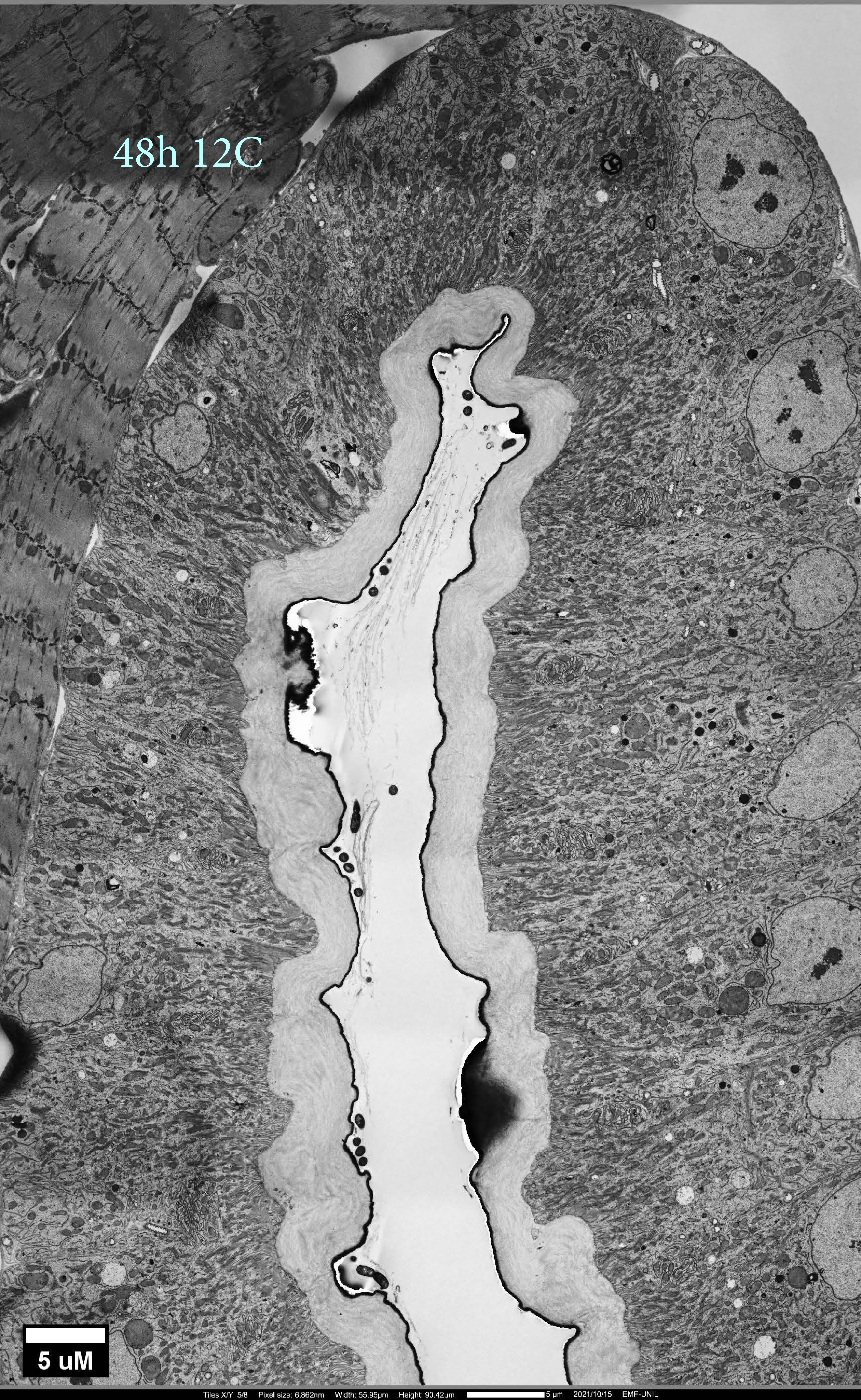

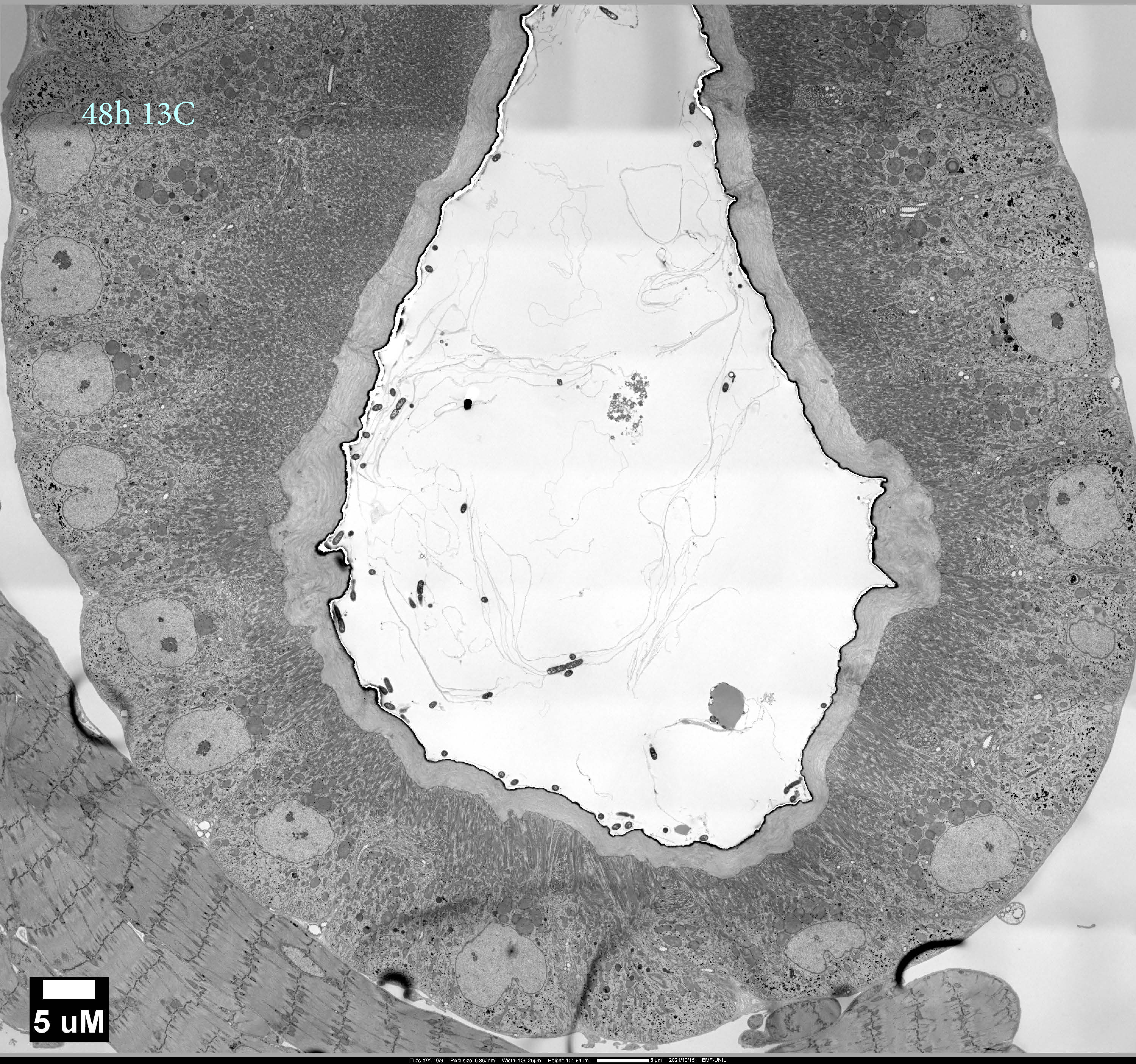

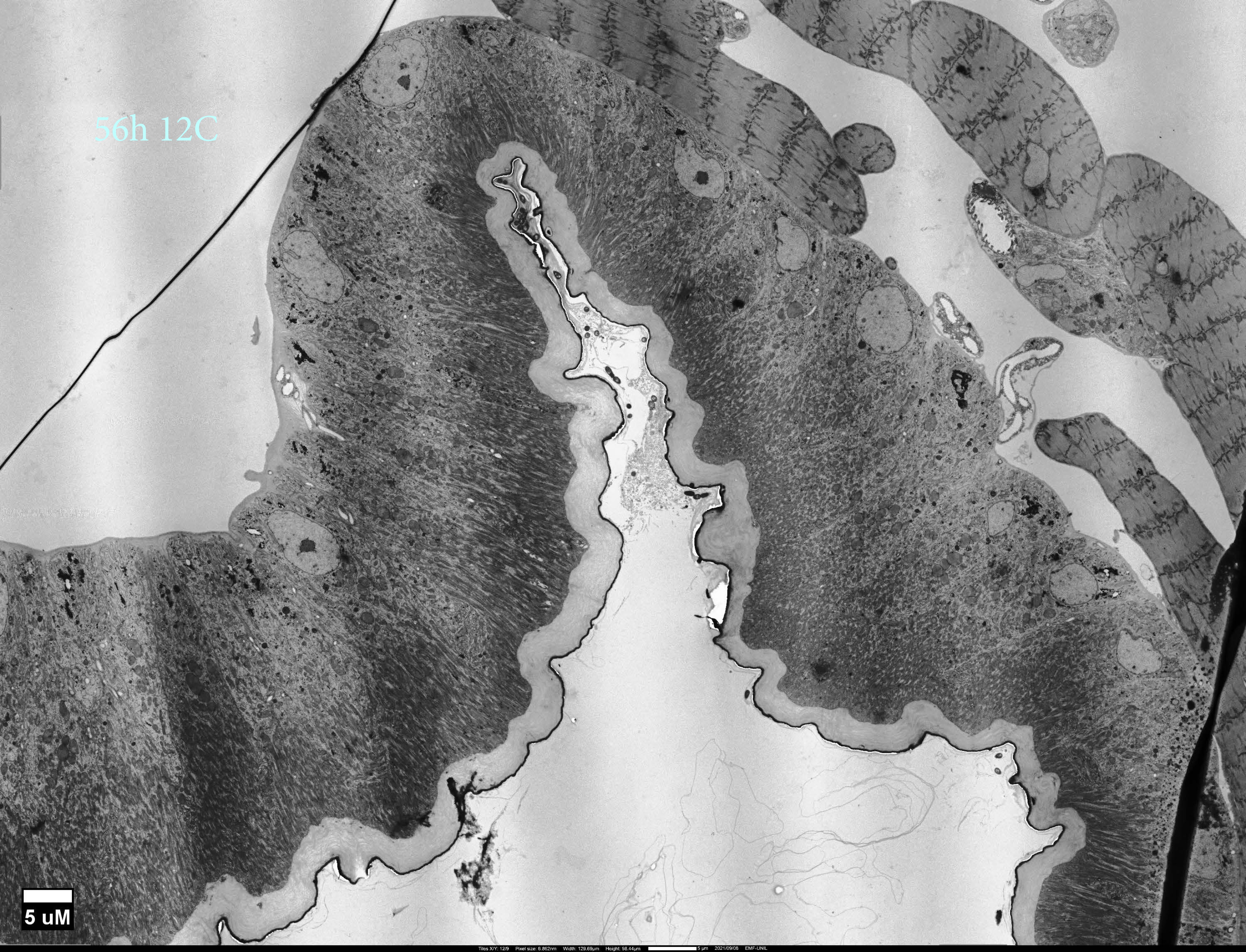

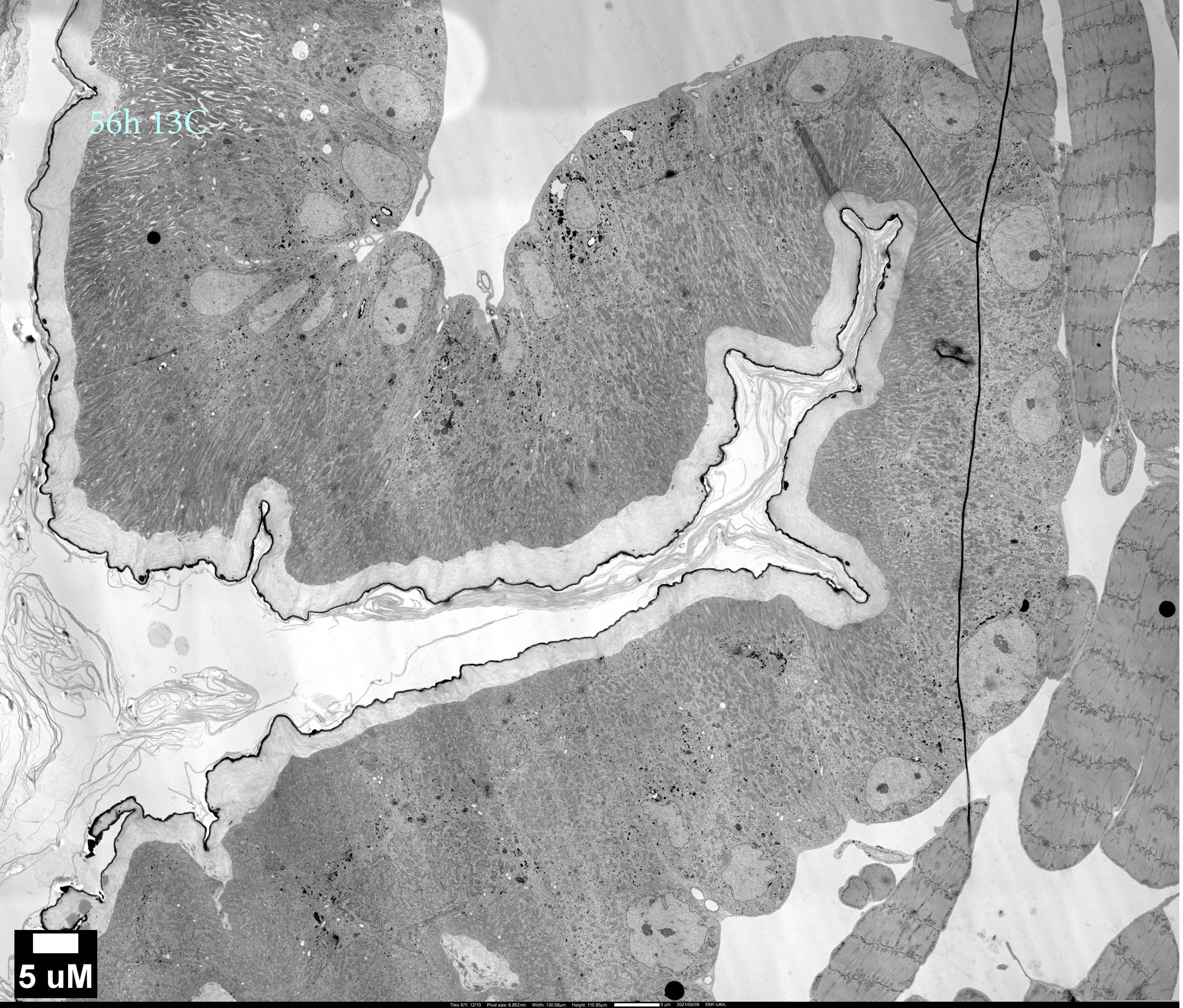

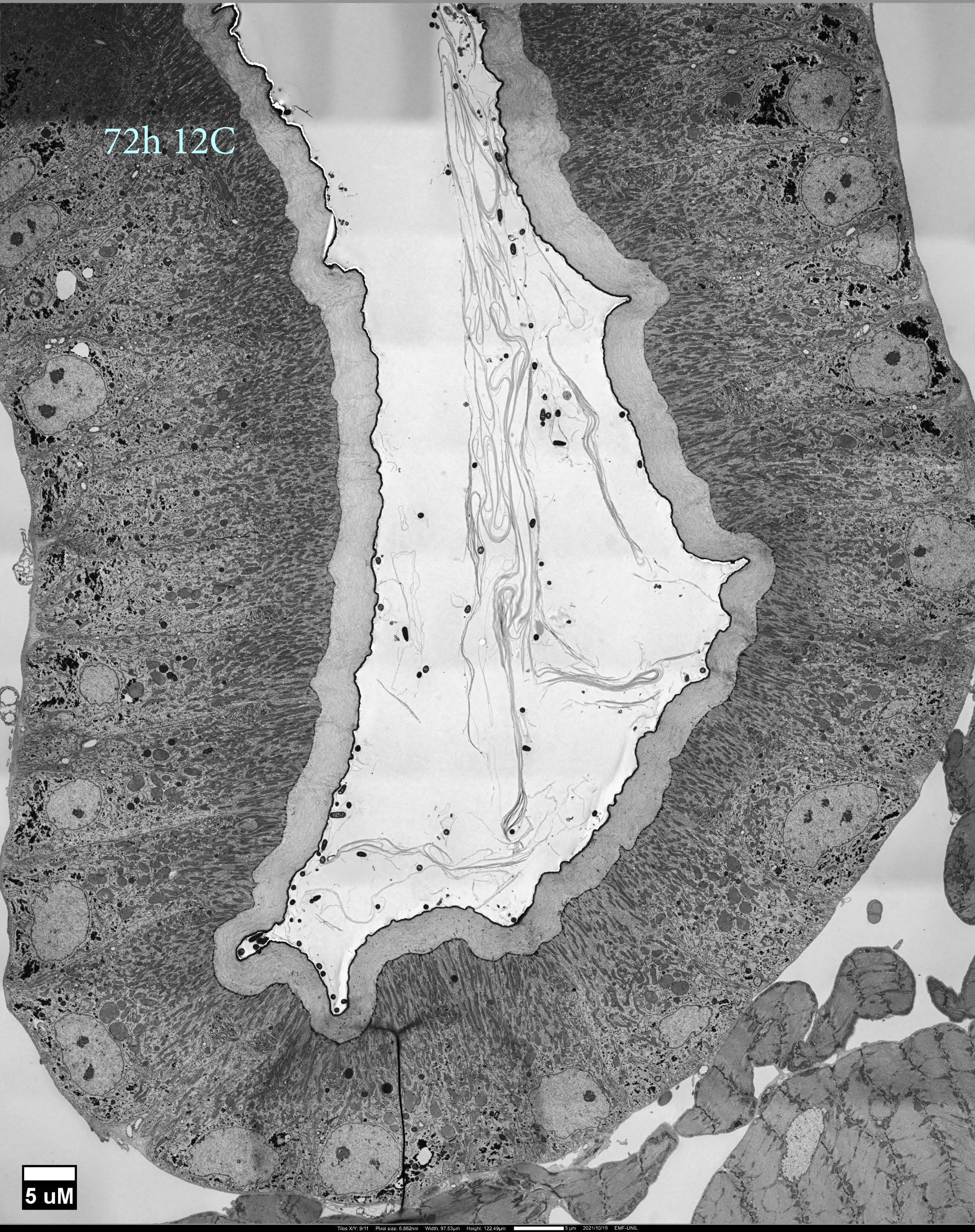

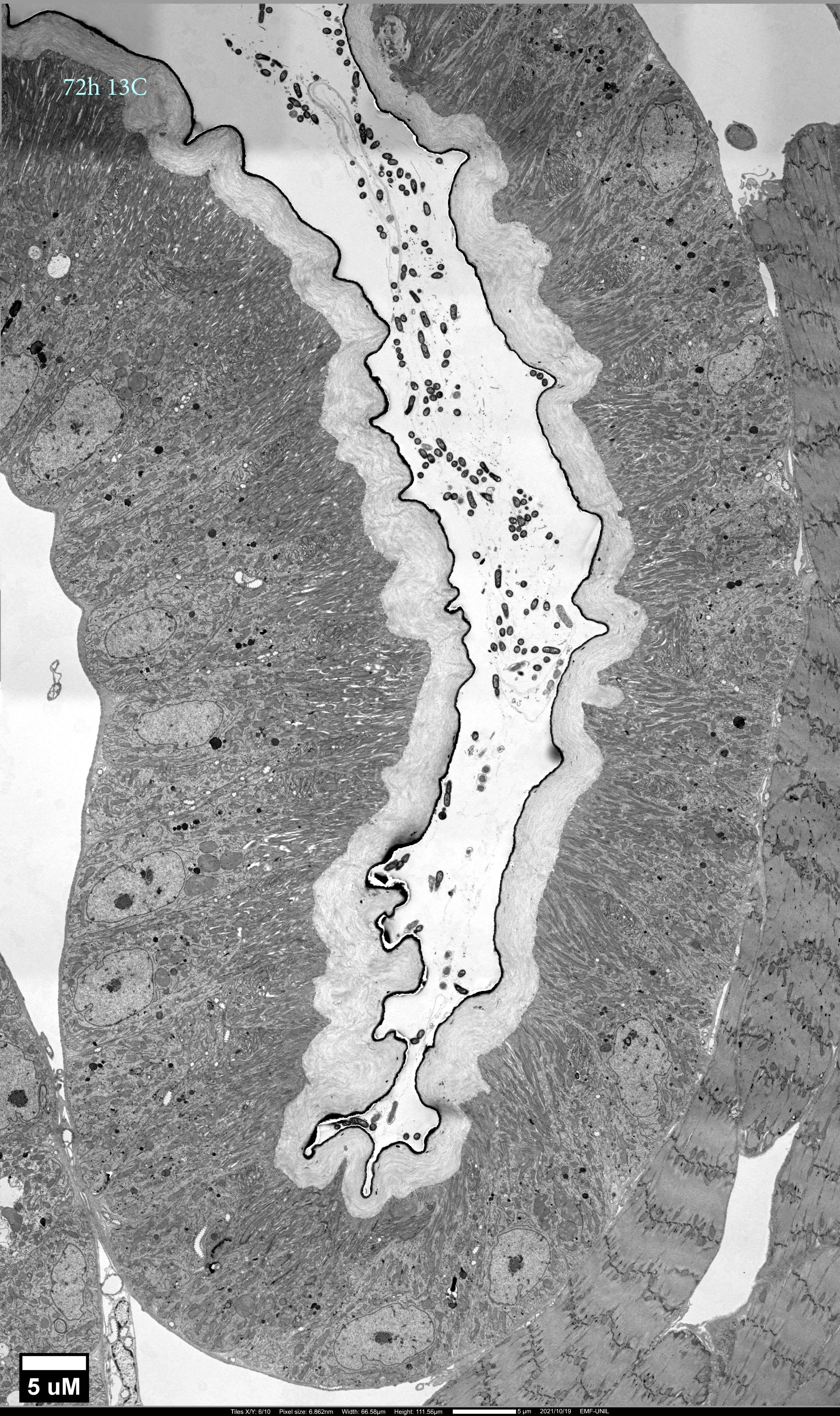
