## Supplementary Dataset 3 for "Foraging on host synthesized metabolites enables the bacterial symbiont *Snodgrassella alvi* to colonize the honey bee gut"

### Bee9: An M9 derived medium for bee gut bacteria

#### Description

This protocol is used to prepare stock solutions for an M9-based 'minimal' medium containing trace metals and vitamins, without casamino acids. Additional vitamins, and growth nutrients are added for growth of *S. alvi* and/or *Gilliamella* strains. The pH should be adjusted to 5.5

---

##### 1. Preparation of HMB (hutner's mineral base) modified metals 44 (component 1)

- Weight on a balance the following:
  1. 1.095g $ZnSO_4$ x $7H_2O$, dissolve in **5ml** $ddH_2O$ at pH=2
  2. 0.914g $FeSO_4$ x $7H_2O$, dissolve in **5ml** $ddH_2O$ at pH=2
  3. 0.154g $MnSO_4$ x $H_2O$, dissolve in **5ml** $ddH_2O$ at pH=2
  4. 0.392g $CuSO_4$ x $5H_2O$, dissolve in **100ml** $ddH_2O$ at pH=2
  5. 0.248g $Co(NO_3)_2$ x $6H_2O$, dissolve in **100ml** $ddH_2O$ at pH=2
  6. 0.177g $Na_2B_4O_7$ x $10H_2O$, dissolve in **100ml** $ddH_2O$ at pH=2
- Mix **1.**, **2.**, **3.**, and add 10ml of each of **4.**, **5.**, **6**. to obtain **component 1**.
- Adjust volume to 100ml and sterile-filter with 0.22 um filter
- Keep it at 4C, light protected

##### 2. Preparation of additional salts (components 2, 3, 4, 5)

- **Component 2:** 1.72g of $MgSO_4$ x $7H_2O$, dissolve in **100ml** $ddH_2O$ at pH=2, sterile-filter with 0.22 um filter
- **Component 3:** 3.33g of $CaCl_2$ x $2H_2O$, dissolve in **100ml** $ddH_2O$ at pH=2, sterile-filter with 0.22 um filter
- **Component 4:** 0.99g of $FeSO_4$ x $7H_2O$, dissolve in **100ml** $ddH_2O$ at pH=2, sterile-filter with 0.22 um filter
- **Component 5:** 0.974g of $(NH_4)_6Mo_7O_{24}$ x $4H_2O$, dissolve in **100ml** $ddH_2O$ at pH=2, sterile-filter with 0.22 um filter
- Keep at RT, light protected

##### 3. Preparation of the vitamin 10x stock solution (component 6a)

- 0.2g of Calcium pantothenate, dissolve in **10ml** $ddH_2O$
- 0.01g of Thiamine-HCl, dissolve in **10ml** $ddH_2O$
- 0.01g of biotin, dissolve in **10ml** $ddH_2O$
- Mix 79ml of $ddH_2O$, 10ml of **2.**, 10ml of **3.**, and 1ml of **4.**
- Sterile-filter the final 100 ml of vitamin stock solution (component 6a)
- Component 6a can be kept for at least 6 months at 4°C in the fridge, protected from light

##### 5. Preparation of 10x stock solution of M9 (component 8)

- Mix the following:
- 60g $Na_2HPO_4$
- 30g $KH_2PO_4$
- 5g NaCl
- 10g $NH_4Cl$
- Add water to 1l, autoclave
- Keep at RT

#### 6. Preparation of *S. alvi* specific vitamins

- Weigh out powder of the following compounds and dissolve in M9 base to a concentration of 10 mM (for final concentration of [0.1 mM]
- 4-Hydroxybenzoate
- P-aminobenzoic acid
- Pyridoxine*HCl
- Choline

#### 7. Prepare 500ml of M9 base medium

- Prepare the final M9 base medium freshly on the day of use
- For 500ml mix the following:
- Component 1: 500 µl
- Component 2: 8390 µl
- Component 3: 1000 µl
- Component 4: 100 µl
- Component 5: 10 µl (100 uL of 1/10 component 5 for small M9 volume)
- Component 6a (10x Vitamin stock): 50ml
- Component 8 (10x M9 stock): 50ml
- Custom vitamins (10mM stock) 5 mL
- dd$H_2O$: 385 ml

#### 8. Prepare carbon + media

- Dissolve compounds of choice in M9 to a final concentration of 10 mM. Citrate is a good single carbon source. Glucose [10 mM] can also be added for *Gilliamella* growth.
- Common substrates include.
    - Alpha-ketoglutaric acid
    - Fumaric acid
    - Malic acid
    - Succinic acid
    - Citric acid
    - Sodium pyruvate
    - Sodium acetate
    - Sodium lactate
  - Check pH and adjust to 5.5 to 6
